# A Thiopurine-like Mutagenic Process Defines TGCT Subtypes

**DOI:** 10.1101/2025.06.12.655573

**Authors:** Kevin M. Brown, Jun Zhong, Adriana Morales Miranda, Mengyan Zhang, Joycelyn Williams, Jacob Williams, Haoyu Zhang, Cheng Liang, Wenbo Li, Bin Zhu, Stephen J. Chanock, Katherine L. Nathanson, Tongwu Zhang

## Abstract

Testicular germ cell tumors (TGCTs) are the most common malignancy in young men, exhibit a unique developmental origin and exceptional chemosensitivity. However, the molecular distinctions between TGCT subtypes remain poorly understood. Here we present a comprehensive genomic analysis of 252 treatment-naive primary TGCTs, integrating deep whole-genome sequencing with matched transcriptomic and epigenomic data. We identify new driver genes and uncover defining features of TGCTs, including pervasive chromosome X amplification with subtype-specific X chromosome inactivation, and a germ cell-like transcriptional program. Although previously reported, whole genome doubling (WGD) in TGCTs is further characterized here as ubiquitous, developmentally early, and associated with age at onset. Seminomas are enriched for early driver mutations, secondary WGD events, sustained *XIST* expression and replication stress-associated indel mutational signatures, while non-seminomas show greater structural complexity, subclonal diversity, relatively earlier-onset WGD, extended tumor latency, and telomere elongation. Moreover, we identify a mutational signature, SBS87, that is exceptionally rare across cancers with exception of thiopurine-treated leukemia, but strikingly prevalent in TGCT, especially non-seminomas. SBS87 is linked to extended tumor latency and telomere elongation, implicating possible environmental or endogenous processes that mimic thiopurine-induced DNA damage in TGCT pathogenesis. Collectively, our findings define TGCTs as molecularly distinct tumors shaped by early genomic instability and highlight SBS87 as a novel mutational footprint with potential etiologic and clinical relevance.

## Introduction

Type II testicular germ cell tumors (TGCTs) are the most common malignancy in young men, predominantly affecting individuals aged 15 to 40 years. TGCTs represent a unique cancer paradigm due to their extreme chemotherapy sensitivity^1^, distinctive histopathology^2,3^, distinctive genomic landscape^4,5^, global DNA hypomethylation^5,6^, and developmental origin from primordial germ cells^7,8^. They are histologically classified into seminomas, which resemble germ cells differentiating along the spermatogenic lineage^9^, and non-seminomas, a heterogeneous group with embryonal differentiation that includes embryonal carcinoma, yolk sac tumors, and teratomas^10^. These subtypes exhibit distinct clinical, histopathological, and molecular characteristics. Seminomas typically present later in young adulthood, demonstrating uniform morphology, marked chemosensitivity, and excellent prognosis^11^. In contrast, non-seminomas arise earlier, display diverse differentiation patterns, exhibit more aggressive behavior with higher metastatic potential, and are associated with a comparatively poorer prognosis^5,12^. Although TGCTs have exceptionally high survival rates due to their sensitivity to platinum-based chemotherapy, treatment-related toxicities and resistance in refractory or recurrent tumors underscore the need for a deeper understanding of their molecular and genomic characteristics^13–15^.

Genomic studies have revealed key alterations driving TGCT pathogenesis. Targeted and exome sequencing studies have identified KIT/RAS pathway mutations, particularly enriched in seminomas^5,6,16^. Additionally, chromosome 12p amplification, a hallmark event in TGCTs, is observed across both subtypes^16^. Insights from The Cancer Genome Atlas (TCGA) based on whole-exome sequencing (WES) have further characterized TGCT genomics, highlighting distinct mutational landscapes, a remarkably low somatic mutation burden compared to other solid tumors, and epigenetic (DNA methylation) differences between seminomas and non-seminomas^5,17^. However, a comprehensive understanding of complex genomic events, genomic instability, mutational signatures, and clonal evolution patterns distinguishing these subtypes remains limited, requiring whole-genome sequencing (WGS) data for analysis. Understanding these genomic distinctions and their interplay is crucial for elucidating tumorigenesis mechanisms and identifying novel therapeutic targets. The recent TGCT WGS dataset (*e.g.*, the Genomics England 100,000 Genomes Project), comprising a relatively small number of patients (55 primary TGCTs)^18–20^, lacked sufficient statistical power to fully address these genomic complexities.

Leveraging comprehensive WGS data from 252 TGCT samples and integrating with RNA sequencing (RNA-Seq) and DNA methylation data from TCGA, we reveal a multifaceted genomic landscape shaped by chromosome X amplification, mutagenesis, and early whole-genome doubling (WGD). Our analysis reveals striking differences between seminomas and non-seminomas in mutagenic processes, X chromosome inactivation (XCI), replication stress, telomere dynamics, and evolutionary patterns. Notably, mutational signature SBS87, linked to prior thiopurine treatment, was identified exclusively in non-seminomas and was significantly associated with tumour latency and telomere length. These findings provide critical insights into the molecular etiology of TGCTs, delineating subtype-specific tumor biology and evolutionary trajectories.

## Results

### Genomic Landscape of Seminomas and Non-Seminomas

We analyzed WGS data from 252 treatment-naive primary TGCT patients from the TCGA study, with a median sequencing depth of 82× for tumor samples and 32× for matched blood-derived normal controls. A subset of patients also had matched RNA-Seq (N=136) and DNA methylation (N=140) data. The cohort included 143 seminomas and 109 non-seminomas, with genetic ancestry estimation based on WGS data identifying 186 patients who clustered with European reference samples (self-reported non-Hispanic White or missing race and ethnicity data) and 66 who did not (including self-reported Black, Asian and Hispanic patients; **Supplementary Data 1**). Non-seminoma subtypes included mixed germ cell tumors (N=55), embryonal carcinomas, non-specific (N=25), benign or malignant teratoma (N=7), yolk sac tumors (N=3), other tumors without subtype information (N=19). Consistent with previous reports, non-seminomas were diagnosed at significantly younger ages than seminomas (median age: 30 vs. 34 years; Wilcoxon test *P*=2.9e-08; **Supplementary Fig. 1**).

Consistent with previous studies, the TCGA TGCT cohort exhibited a relatively low number of somatic alterations, likely reflecting their unique embryological origin^5,16,20^. The median tumor mutational burden (TMB) was 0.45 mutations/Mb (range: 0.04–1.62), the median percentage of the genome altered (PGA) by copy number aberrations was 85.9% (range: 0.7%–95.8%), with median 27.5 structural variants (SVs) (range: 0–154) and median 0 transposable element insertions (TEs) (range: 0–20; mean: 0.84). Notably, non-seminomas exhibited significantly higher levels of genomic alterations for TMB, PGA, SVs, and TEs, compared to seminomas (FDR-adjusted *P* values from linear regression, corrected for age and tumor purity: TMB=0.049; PGA=7.11e-04; SVs=1.24e-06; TEs=0.005; **Fig. 1a**). Genetic ancestry was not significantly associated with any of the somatic alteration metrics (FDR > 0.05 for all comparisons).

**Fig. 1:**
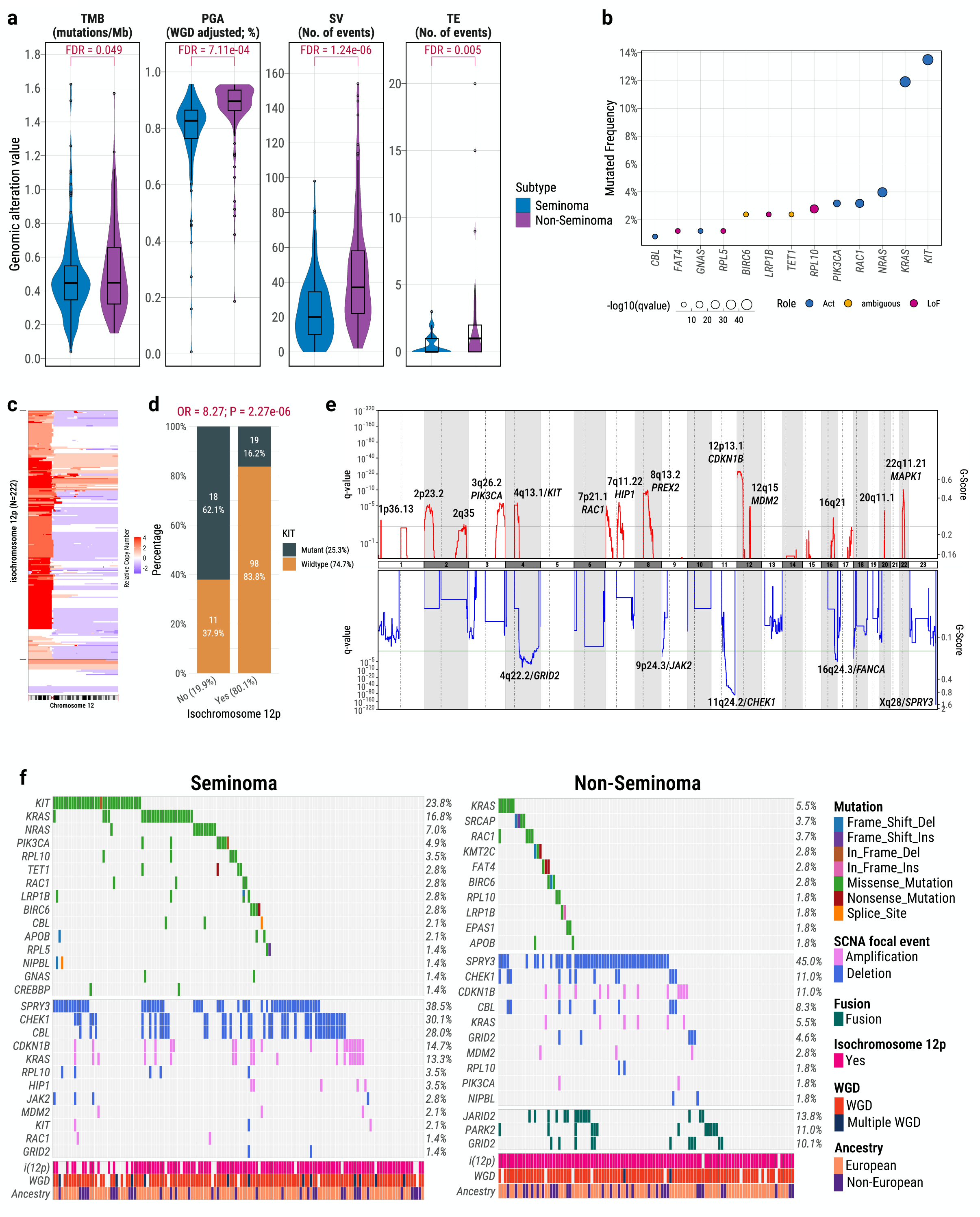
Distinct Genomic Landscapes of Seminomas and Non-Seminomas in Testicular Germ Cell Tumors (TGCTs). (**a**), Comparison of key genomic features between seminomas (blue) and non-seminomas (purple), including tumor mutational burden (TMB; mutations/Mb), percentage genome altered (PGA; WGD-adjusted), number of structural variants (SVs), and number of transposable element insertions (TEs). P-values are from linear regression models adjusted for age and tumor purity; FDR correction via Benjamini–Hochberg method. (**b**), Significantly mutated genes identified by the IntOGen pipeline across all TCGT samples. Dot size reflects combined -log10 q-value based on different driver gene algorithms. Colors denote functional annotation (Act: activating; LoF: loss-of-function). (**c**), Copy number heatmap of chromosome 12 showing frequent 12p amplification and isochromosome 12p [i(12p)] in TGCTs. (**d**), Enrichment of i(12p) in seminomas with *KIT* mutations. P-value and odds ratio from two-sided Fisher’s exact test shown above the barplot. (**e**), GISTIC2.0 analysis of recurrent focal copy number alterations. Significant amplifications (red) and deletions (blue) are annotated with potential target genes. Y-axis represents the significance by q value (left) and normalized amplification signals (G-Score; right). The green line represents the significance cutoff at Q value=0.25. (**f**), Oncoprints depicting recurrent driver mutations, focal amplifications/deletions, and gene fusions in seminomas and non-seminomas. Mutation frequencies for each gene are shown on the right. For mutations, only recurrent nonsynonymous mutations in known pan-cancer driver genes were included. For focal somatic copy number alterations (SCNAs), only high-level amplifications (copy number >= 2) and deep deletions (copy number = –2) were considered. For gene fusions, we included only recurrent events involving known pan-cancer driver genes. All boxplots display median, interquartile range (IQR), and whiskers extending to 1.5× IQR.

### Subtype-Specific Drivers and Structural Complexity in TGCTs

Using the IntOGen pipeline^21^, which integrates seven complementary driver-discovery algorithms, we identified 13 significant TCGT driver genes (**Fig. 1b**), collectively mutated in 58.0% of seminomas and 19.3% of non-seminomas, highlighting greater genetic heterogeneity in non-seminomas. The majority (85%) of these driver mutations were clonal, suggesting early occurrence during tumor evolution (**Supplementary Fig. 2**; **Supplementary Data 2**). Consistent with previous studies, the most frequently mutated driver genes in TGCTs involved the RAS signaling pathway^22^, including *KIT*, *KRAS*, *RAC1*, and *PIK3CA*, predominantly occurring in seminomas, as well as *NRAS* and *CBL*, whose mutations occurred exclusively in seminomas. *KIT* mutations predominantly clustered in activating hotspots (exons 11 and 17)^23^, while *KRAS* mutations frequently affected activating codons, with G12 (n=15), Q61 (n=5), and A146 (n=3) observed in seminomas, and G12 (n=4), Q61 (n=1), and A146 (n=1) in non-seminomas (**Supplementary Fig. 3**). Additionally, we identified rare driver mutations (<3%) in genes not previously reported in TGCTs, including *FAT4*, *GNAS*, *RPL5*, *BIRC6*, *LRP1B*, *TET1*, and *RPL10*, most of which are involved in cell growth and proliferation pathways. Notably, *TP53* mutations were entirely absent, consistent with previous observations in TGCT and other germ-cell tumors, reflecting their unique primordial germ cell origin^6,24^.

Allele-specific copy number alteration analyses identified frequent arm-level copy number gain involving chromosomes 12p (88.9%), 21 (32.9%), 7 (28.2%), 8 (23.0%), X (20%), and 12q (15.5%). Tumors exhibiting co-occurrence of these amplifications clustered together (N=36; 14.3%; cluster 3 in **Supplementary Fig. 4**) and were significantly depleted for samples with whole-genome doubling (WGD; 44.4% with WGD) relative to the samples in two distinct clusters (99.5% WGD; Fisher’s exact test, *P*=7.1e-19, odds ratio/OR=0.004; **Supplementary Fig. 4**). Isochromosome 12p [i(12p)], a hallmark of germ cell tumors characterized by amplification of chromosome arm 12p^6,16,25,26^, was detected in 88% of TGCTs (**Fig. 1c**), with significantly higher frequency in non-seminomas (98%) compared to seminomas (80%; Fisher’s exact test, *P* = 6.11e-06), consistent with previous reports^5,27^. Only two non-seminomas lacked i(12p). Among seminomas, tumors without i(12p) were significantly enriched for *KIT* mutations (Fisher’s exact test, *P*=2.27e-06; **Fig. 1d**). Applying the GISTIC algorithm to all TCGT tumors^28^, we identified known recurrent focal copy number alterations, including 13 significant amplifications containing known or potential driver genes (e.g., 12p13.1/*CDKN1B*, 12q15/*MDM2*, 4q13.1/*KIT*, 22q11.21/*MAPK1*) and five significant deletions (e.g., 4q22.2/*GRID2*, 11q24.2/*CHEK1*, 16q24.3/*FANCA*, 9p24.3/*JAK2*) (**Fig. 1e)**. Most of these focal events occurred in both subtypes but were more significantly targeted in seminomas, except 9p24.3/*JAK2* deletion, exclusively found in seminomas (**Supplementary Fig. 5**). For example, seminomas showed significantly higher frequency of 11q24.2 (*CHEK1*/*CBL*) deletions compared to non-seminomas (30% *vs.* 11%; Fisher’s exact test, *P* = 3.41e-04). Notably, the most significant and common focal alteration targeted the pseudoautosomal region (PAR) at Xq28 in both subtypes (deleted in 38.5% of seminomas; 45% of non-seminomas; **Fig.1e**), involving deletion of *SPRY3*, a potential negative regulator of the RAS/MAPK signaling pathway. This alteration is particularly notable given that *SPRY4*, a paralog of *SPRY3* and a major TGCT GWAS susceptibility locus^29,30^, also modulates the same pathway. These findings raise the possibility of a shared germline–somatic axis in TGCT pathogenesis involving SPRY-mediated RAS/MAPK regulation, and point to a potential tumor-suppressive role for *SPRY3* that warrants further investigation. To characterize the architecture of focal amplifications, we applied AmpliconArchitect^31^, and classified amplification events into linear (17.9% of all TGCT samples), extrachromosomal DNA (ecDNA; 6.8%), breakage-fusion-bridge (BFB) cycles (1.6%), or other complex rearrangements (4.4%). As expected, ecDNA events in TGCTs occur at a lower frequency (5.6% in seminomas and 8.3% in non-seminomas) compared to most other solid tumors^32^ and exhibit the highest copy-number amplifications (**Supplementary Fig. 6a**). Notably, ecDNA contributed to amplification of oncogenes including *KRAS* (5 seminomas), *KIT* (1 seminoma), and *MDM2* (1 seminoma, 2 non-seminomas) (**Supplementary Fig. 6b-e)**.

Moreover, we identified several recurrent gene fusions, exclusively in non-seminomas (**Fig. 1f**). These fusions included *JARID2* (13.8%), a component of the Polycomb Repressive Complex 2; *PARK2* (11.0%), a regulator of the ubiquitin-proteasome system; and *GRID2* (10.1%), which is involved in glutamatergic signaling. No recurrent gene fusions were observed in seminomas.

In summary, seminomas exhibited more driver mutations (overall 58% vs. 19.3%; Fisher’s exact test; P=3.59e-10) and focal amplifications. Conversely, non-seminomas showed extensive SVs and fusion events, reflecting their greater genomic complexity and intratumoral heterogeneity.

### Frequent Chromosome X Amplification and Inactivation in TGCTs

Consistent with prior reports^5,16^, our WGS data analysis revealed extensive autosomal chromosome losses accompanied by gains in chromosome X (**Supplementary Fig. 4**). After adjusting for tumor purity, nearly all TGCT samples showed significantly higher chromosome X-to-autosome sequencing depth ratios compared to matched blood (Wilcoxon paired test, *P*=1.06e-42), indicating widespread chromosome X amplification (**Fig. 2a**). Non-seminomas exhibited significantly higher chromosome X amplification levels than seminomas (Wilcoxon test, *P*=1.37e-12). Absolute and subclonal copy-number analyses further confirmed chromosome X gain spanning at least 50% of chromosome X in 94% of TGCTs (96.3% in non-seminomas, 91.6% in seminomas) (**Fig. 2b**; **Supplementary Data 3**); up to eight copies of chromosome X were observed in these samples. Additionally, subclonal chromosome X copy-number gain occurred in 17.7% of tumors (15.4% seminomas, 20.2% non-seminomas), highlighting the dynamic nature of chromosome X amplification in TGCTs.

**Fig. 2:**
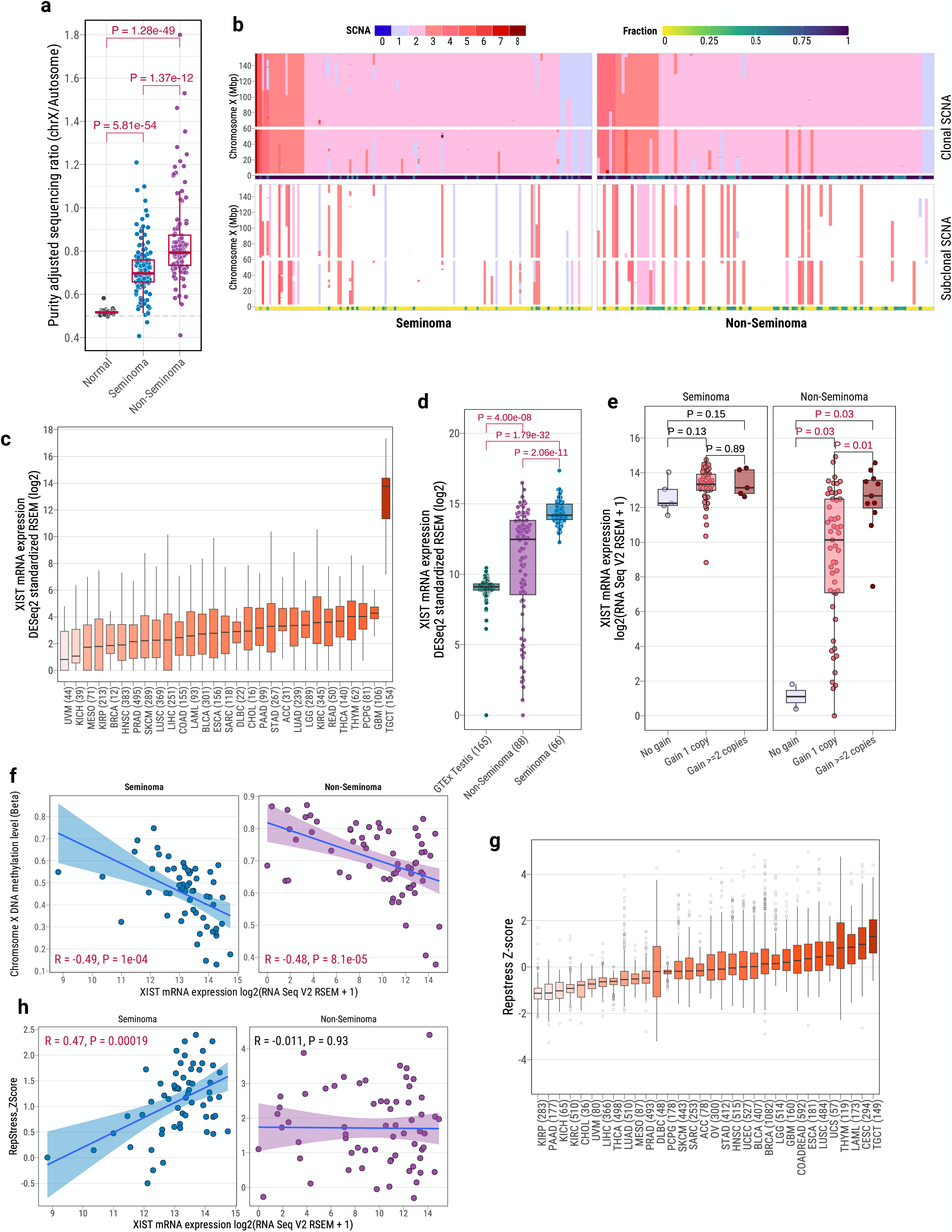
Widespread Chromosome X Amplification and X Chromosome Inactivation (XCI) in TGCTs. (**a**) Tumor purity-adjusted sequencing depth ratio of chromosome X versus autosomes in matched normal blood, seminomas, and non-seminomas. P-values from two-sided Wilcoxon rank-sum tests are shown above boxplots. (**b**) Copy number profiles of chromosome X in TGCTs, separated by seminoma (left) and non-seminoma (right), ordered by total copy number. The y-axis indicates chromosome X positions. Top panel: clonal somatic copy number alterations (SCNAs); bottom panel: subclonal SCNAs. Subclonal fractions per sample are shown in the barplot below each panel. (**c**) *XIST* mRNA expression across TCGA tumor types (male subjects), with TGCTs showing the highest levels. (**d**) *XIST* mRNA expression in normal testis (GTEx), non-seminomas, and seminomas (TCGA). P-values from two-sided Wilcoxon rank-sum tests are shown above boxplots. (**e**) Association between chromosome X copy number and *XIST* expression in seminomas and non-seminomas, with significant correlation in non-seminomas only. *P*-values from two-sided Wilcoxon rank-sum tests are shown above boxplots. (**f**) Negative correlation between *XIST* expression and chromosome X median DNA methylation in seminomas (left) and non-seminomas (right). Pearson correlation coefficient (R) and P-value are shown below. (**g**) Replication stress scores across TCGA tumor types based on replication stress gene signatures, with TGCTs showing the highest levels. (**h**) Correlation between *XIST* expression and replication stress scores in seminomas (left) and non-seminomas (right), with significant positive correlation in seminomas only. Pearson R and P-value are shown above. All boxplots display median, IQR, and whiskers extending to 1.5× IQR.

In contrast, chromosome Y-to-autosome ratios suggested frequent Y chromosome loss across 252 TGCTs (**Supplementary Fig. 7a**). Large arm-level Y deletions (>10 Mbp) were observed in 23.4% of tumors, with similar frequencies in seminomas (25.9%) and non-seminomas (20.2%) (Fisher’s exact test; P = 0.37; **Supplementary Fig. 7b**). Additionally, focal deletions targeting the RBMY gene family (*RBMY1A1*, *RBMY1B*, *RBMY1D*)—male germ cell–specific RNA-binding proteins—were significantly enriched in seminomas (26.6% vs. 12.8% in non-seminomas; P = 0.0078). Notably, 6.3% of TGCTs showed copy number gains on chromosome Y, with a significant enrichment in non-seminomas (11.0% vs. 2.8% in seminomas; Fisher’s exact test; P = 0.016). A recurrent copy number gain was also observed across different regions of Yp11.2, encompassing the TSPY gene family.

Given the prevalence of chromosome X amplification, we explored potential X chromosome inactivation (XCI), a process in female mammals to epigenetically silence one of the two X chromosomes to maintain balanced gene dosage between sexes^33,34^. We hypothesized that TGCTs may initiate XCI in response to multiple chromosome X copies. To evaluate this, we examined expression of the long non-coding RNA *XIST* (X-inactive specific transcript), a known marker and key regulator of XCI^35–37^. Strikingly, compared to other tumor types from male patients across TCGA, TGCTs exhibited the highest *XIST* mRNA expression levels (**Fig. 2c**). Consistent with this observation, TGCTs also showed significantly elevated *XIST* expression compared to normal testis tissue from the Genotype-Tissue Expression (GTEx) study (fold change=25.2; **Fig. 2d**). Seminomas consistently exhibited elevated *XIST* expression, while non-seminomas showed greater variability (Seminomas *vs.* non-seminomas, linear regression adjusted tumor purity, *P*=3.81e-07). Notably, a significant association between the number of chromosome X copies and *XIST* expression was detected specifically in non-seminomas (linear regression adjusted tumor purity, *P*=4.63e-04; **Fig. 2e**), but not in seminomas (*P*=0.14). This difference may reflect subtype-specific strategies for managing extra X chromosome copies, with seminomas potentially relying more on constitutive *XIST*-driven for chromosome X silencing, whereas non-seminomas may modulate *XIST* expression according to the extent of chromosome X amplification.

XCI is typically accompanied by distinct epigenetic changes, including histone acetylation and DNA methylation, to stabilize gene silencing^38^. Given the global DNA hypomethylation characteristic of TGCTs, especially seminomas^5,39^, we next compared DNA methylation levels between chromosome X and autosomes. In both TGCT subtypes, chromosome X exhibited lower methylation levels compared to autosomes (**Supplementary Fig. 8**). A significant negative correlation was observed between *XIST* expression and DNA methylation levels on chromosome X in both subtypes (Pearson correlation; seminomas: *R* = -0.49, *P* = 1e-04; non-seminomas: *R* = -0.48, *P* = 8.1e-05; **Fig. 2f**). This reduced methylation on chromosome X is consistent with observations from XCI studies of female tissues^40–43^, where the X chromosome is globally hypomethylated, suggesting active selection for a hypomethylated state on the X chromosome in TGCTs.

The epigenetic silencing of chromosome X through XCI can lead to replication stress due to tighter chromatin packaging and reduced DNA accessibility during replication^44,45^. To investigate this connection in TCGT, we computed replication stress scores based on the transcriptional profiles of replication stress gene signatures, reflecting cellular characteristics indicative of replication stress^46^. Interestingly, TGCTs exhibited the highest levels of replication stress across all TCGA tumor types. (**Fig. 2g**; **Supplementary Data 4**). After adjusting for tumor purity and age, no significant difference in replication stress was observed between seminomas and non-seminomas (**Supplementary Fig. 9a**). Among the limited number of tumors lacking chromosome X gains, neither seminomas nor non-seminomas showed a significant difference in replication stress compared to those with chromosome X amplification (**Supplementary Fig. 9b**). Furthermore, seminomas exhibited a strong positive correlation between *XIST* expression and replication stress scores (*R* = 0.47, *P* = 1.9e-04), which was absent in non-seminomas (**Fig. 2h**). Additionally, we observed no evidence of gene escape from XCI, suggesting robust epigenetic regulation of X-linked genes in TGCTs (**Supplementary Fig. 10**).

Taken together, these findings reveal distinct strategies for X chromosome dosage compensation between TGCT subtypes. Seminomas predominantly maintain chromosome X silencing through persistent *XIST* expression, reflecting a primordial germ cell-like epigenetic state that directly contributes to elevated replication stress. In contrast, non-seminomas adjust *XIST* expression according to chromosome X amplification, enabling stable XCI and managing replication stress differently. These epigenetic and genomic differences suggest a potential critical role of XCI in determining the distinct histological and developmental trajectories observed in TGCT subtypes.

### Distinct Mutational Processes Define TGCT Subtypes

Although the overall mutational burden was similar between seminomas and non-seminomas, unsupervised hierarchical clustering clearly separated the two TGCT subtypes based on the mutational spectrum (**Fig. 3a**). Leveraging extensive whole-genome sequencing (WGS) data from TGCT samples, we systematically identified operative mutational signatures using SigProfilerExtractor^47^ and robustly quantified their activities through parametric bootstrapping^48^ (**Methods**; **Supplementary Data 5**). We identified five single base substitution (SBS) signatures within TGCT samples (**Fig. 3b**): two clock-like mutation signatures (SBS1 and SBS5; **Supplementary Fig. 11**), one signature associated with defective homologous recombination (HR) DNA repair (SBS3), one indicative of reactive oxygen species (ROS)-related damage (SBS18), and one previously linked to thiopurine treatment (SBS87). Notably, SBS3 and SBS5 were dominant across nearly all 252 TGCT samples, contributing substantially to the mutational load in both subtypes (seminomas: 39.2% SBS3 and 58.3% SBS5; non-seminomas: 30.2% SBS3, and 56.4% SBS5). Similarly, six small insertion and deletion (ID) signatures were identified, including those linked to DNA replication slippage (ID1/ID2), defective HR-based DNA damage repair (ID6), and signatures of unknown etiology (ID4, ID5, ID9) (**Fig. 3c**). Intriguingly, ID signatures (ID1, ID2, and ID9) were significantly associated with DNA replication stress exclusively in seminomas (**Fig. 3d**), consistent with prior reports implicating replication-associated DNA damage in TGCT pathogenesis^49^ .

**Fig. 3:**
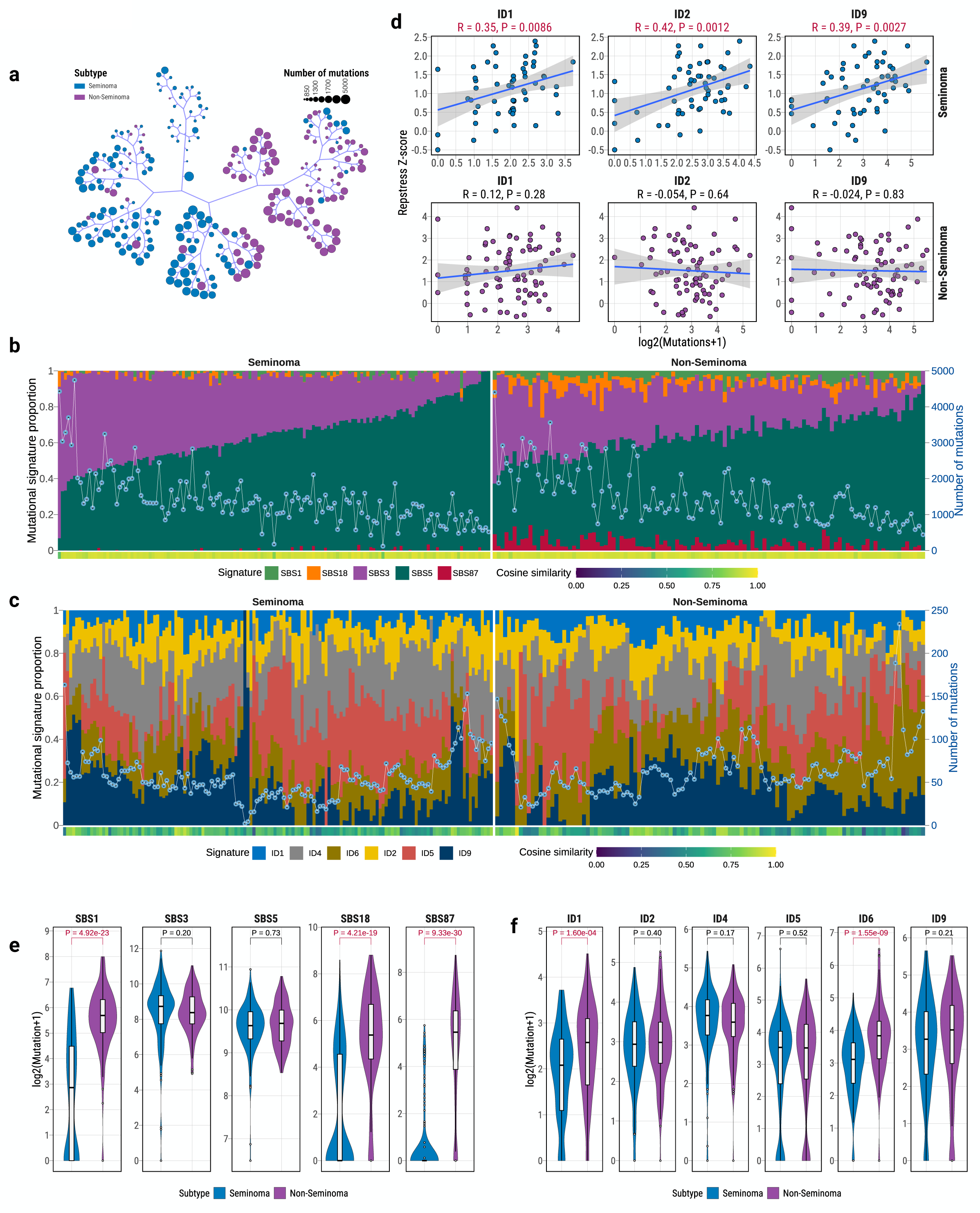
Distinct Mutational Processes in TGCT Subtypes. (**a**) Unsupervised hierarchical clustering of TGCTs based on single-base substitution (SBS) and insertion-deletion (ID) mutational profiles, separating seminomas (blue) and non-seminomas (purple). (**b**) SBS mutational signature decomposition, with stacked barplots showing relative contributions per signature (left y-axis). A line with dots indicates the number of mutations (right y-axis). Cosine similarity between original and decomposed profiles is shown below. (**c**) ID mutational signature decomposition, with stacked barplots showing relative contributions per signature (left y-axis). A line with dots indicates the number of mutations (right y-axis). Cosine similarity between original and decomposed profiles is shown below. (**d**) Correlation between mutational burden of ID signatures (ID1, ID2, ID9) and replication stress scores, with significant positive correlations in seminomas only. Pearson R and P-value are shown above scatterplots. (**e**) Comparison of SBS mutational signature burden between seminomas and non-seminomas. Statistical significance (FDR) was assessed by linear regression, adjusted for tumor purity, with Benjamini–Hochberg correction. (**f**) Comparison of ID mutational signature burden between subtypes, with non-seminomas showing higher ID1 and ID6 activity. Statistical significance (FDR) was assessed by linear regression, adjusted for tumor purity, with Benjamini–Hochberg correction. All boxplots display median, IQR, and whiskers extending to 1.5× IQR.

Homologous recombination deficiency (HRD)-associated signatures (SBS3, ID6) were pervasive across both TGCT subtypes (present in 99.6% and 97.2% of samples, respectively)^18^. This observation aligns with previous evidence that TGCTs exhibit heightened sensitivity to platinum-based chemotherapy due to underlying DNA repair impairments^18,50^. A striking divergence in mutational signature distribution was observed between seminomas and non-seminomas, with non-seminomas showing significant enrichment mutations attributed to signatures SBS1, SBS18, SBS87, ID1, and ID6 (**Fig. 3e-f**). Notably, SBS87 has been previously identified in relapsed acute lymphoblastic leukemia (ALL) and experimentally validated as associated with thiopurine treatment^51–53^, which is widely used in treatment of ALL as well as autoimmune disorders. On the surface, the high prevalence of SBS87 in non-seminomas (86.2%) might suggest prior thiopurine exposure, raising the possibility that thiopurines could also affect other rapidly dividing cell populations beyond leukocytes, such as spermatogonial cells^54,55^. However, given that these samples were treatment-naive and that clinical histories regarding prior immunosuppressive use or diagnosis of other disorders were unavailable, widespread therapeutic exposure is unlikely to account for the ubiquity of this signature in non-seminomas. This raises the possibility that SBS87 may, in part, reflect an endogenous mutagenic process that phenocopies thiopurine-induced DNA damage. In fact, differential expression analysis of 15 key thiopurine metabolism genes^56^ revealed that 80% were significantly dysregulated between subtypes, pointing to potential metabolic differences in thiopurine processing between seminomas and nonseminomas (**Supplementary Fig. 12)**. Taken together, these results underscore distinct mutational processes that shape the genomic landscapes of seminomas and non-seminomas, providing insights into the etiological mechanisms underlying TGCT development and progression.

### Impact of Mutational Signature SBS87 on Tumor Latency and Telomere Length

TGCT subtypes exhibit significant differences in age at diagnosis, making it essential to investigate how distinct genomic alterations and mutational processes influence TGCT age of onset or tumor latency. Tumor latency is defined as the time between the appearance of the most recent common ancestor (MRCA) and the age at diagnosis. Understanding tumor latency is crucial for improving early detection and therapeutic strategies, as shorter latency often indicates aggressive but clonal tumors that are easier to target, while longer latency may lead to greater subclonal diversity, complicating treatment. Using stably accumulating clock-like mutations, particularly CpG>TpG transitions, we estimated MRCA ages and tumor latency, as commonly applied in other studies^57–59^. Consistent with their earlier age at diagnosis, non-seminomas had significantly lower MRCA ages (median: 17.4 years) compared to seminomas (median: 34.3 years; Wilcoxon test, *P*=7.65e-09; **Fig. 4a**). Surprisingly, non-seminomas exhibited significantly longer latency (9.2 years) than seminomas (4.9 years; *P* = 0.0045). These findings suggest that non-seminomas may present earlier but provide a larger window for early detection despite their aggressive nature. Next, we explored genomic factors potentially influencing tumor latency. Genomic alterations in major driver genes such as *KIT* and *KRAS* were not significantly associated with latency (**Supplementary Fig. 13**). However, when examining mutational signatures, we identified a significant positive correlation between the SBS87 signature and tumor latency in both seminomas (Spearman correlation; *R*=0.45, *P*=0.0024) and non-seminomas (*R*=0.27, *P*=0.022; **Fig. 4b-c**). A similar correlation was observed for SBS5 (**Supplementary Fig. 14**), consistent with its known clock-like etiology. These results suggest that the presence of SBS87 may influence tumor evolution and prolong latency, particularly in non-seminomas.

**Fig. 4:**
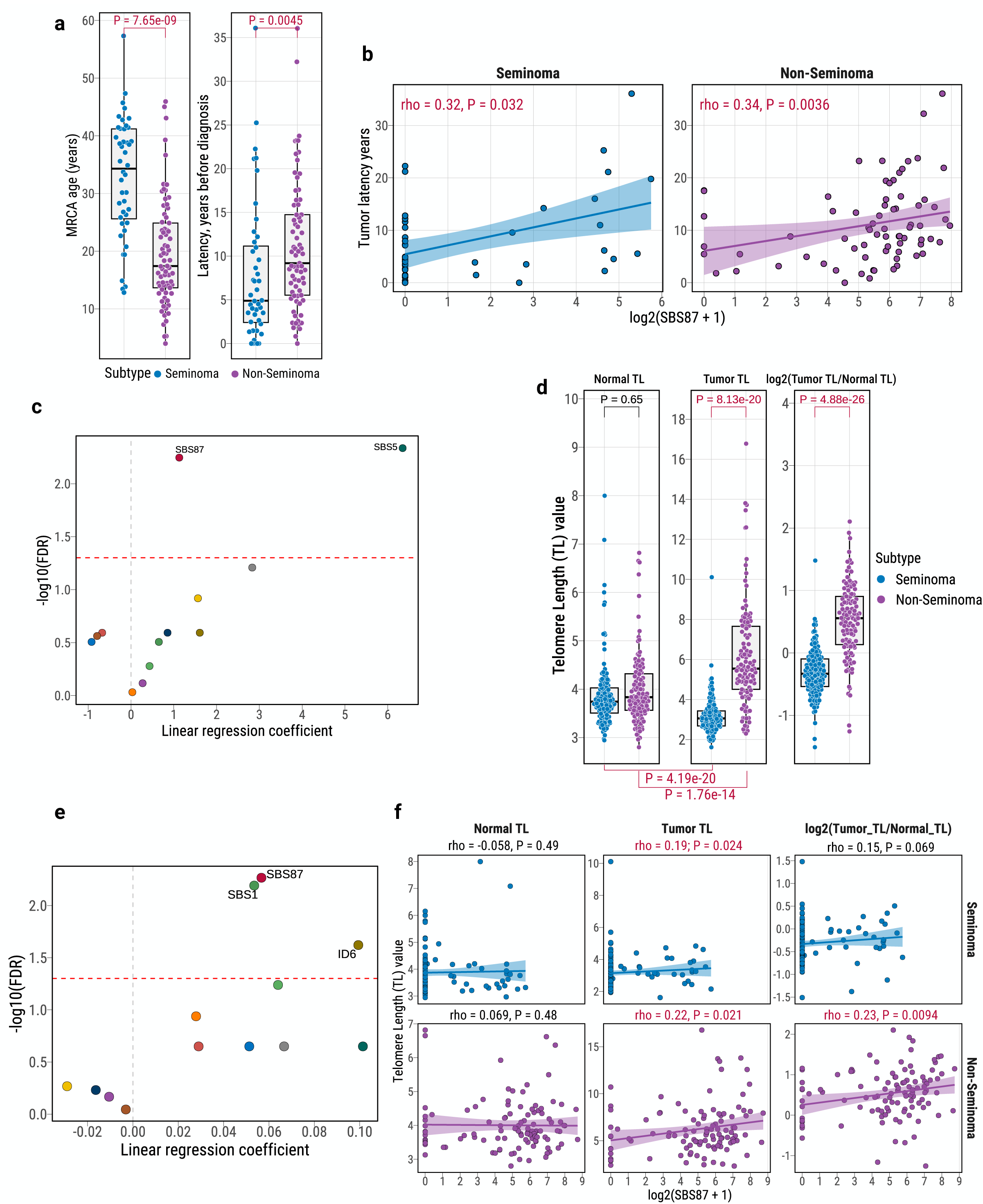
Impact of Thiopurine-Associated Mutational Signature SBS87 on Tumor Latency and Telomere Length in TGCTs. (**a**) Comparison of estimated age of the most recent common ancestor (MRCA, left) and tumor latency (time from MRCA to diagnosis, right) between seminomas and non-seminomas. P-values from two-sided Wilcoxon rank-sum tests are shown above boxplots. (**b**) Correlation between SBS87 activity (log2-transformed) and tumor latency in seminomas (left) and non-seminomas (right). Pearson R and P-value are shown above scatterplots. (**c**) Linear regression analysis of mutational signatures associated with tumor latency in all TGCT samples, adjusted for tumor purity. FDR from Benjamini–Hochberg correction is shown, with FDR<0.05 thresholds marked by a red dashed line. (**d**) Comparison of telomere length (TL) in normal blood, tumor, and tumor/normal TL ratio (log2) between seminomas and non-seminomas. P-values above boxplots are from linear regression, adjusted for tumor purity and age, comparing seminomas and non-seminomas; P-values below are from two-sided Wilcoxon rank-sum tests comparing tumor versus normal TL. (**e**) Linear regression analysis of mutational signatures associated with tumor TL in non-seminomas, adjusted for tumor purity and age. FDR from Benjamini–Hochberg correction is shown, with FDR<0.05 thresholds marked by a red dashed line. (**f**) Correlation between SBS87 activity and TL in normal tissues (left), tumors (middle), and tumor/normal TL ratio (right), stratified by subtype. Significant positive correlations with tumor TL are observed in non-seminomas only. Pearson R and P-value are shown above scatterplots. All boxplots display median, IQR, and whiskers extending to 1.5× IQR.

Telomeres play a crucial role in genome stability, regulating chromosome movement, and facilitating recombination during cell division^60–62^. Previous TCGA studies using whole-exome sequencing (WES) reported longer telomere lengths (TLs) in TGCTs, with nearly all samples expressing *TERT*^63^. However, the molecular mechanisms underlying telomere length regulation in TGCTs remain unclear. Using WGS data, we estimated telomere lengths in both TGCT subtypes. Seminomas exhibited significantly shorter telomeres compared to matched normal tissues (Wilcoxon test, *P*=4.19e-20), while non-seminomas showed significantly longer telomeres (*P*=1.76e-14; **Fig. 4d**; **Supplementary Data 6**). Telomere shortening was observed in 82.5% of seminomas, whereas 80.7% of non-seminomas displayed telomere elongation, aligning with previous reports^64^. To investigate potential genomic factors influencing telomere length, we assessed associations with major driver genes and mutational signatures. No significant association was found between driver gene mutations and tumor telomere length (**Supplementary Fig. 15**). However, SBS87 was significantly associated with telomere elongation in non-seminomas (*FDR* = 0.0054; **Fig. 4e-f**). In summary, our findings indicate that the thiopurine-associated mutational signature SBS87 is significantly correlated with longer tumor latency and telomere elongation in non-seminomas. This suggests a potential role for SBS87 in influencing tumor evolution and genomic stability, providing valuable insights into TGCT progression and identifying possible therapeutic vulnerabilities.

### Divergent Clonal Dynamics Between Seminomas and Non-Seminomas

Given the striking differences in mutational processes and genomic alterations between seminomas and non-seminomas, we next investigated how these features shape clonal evolution and intratumoral heterogeneity in TGCTs. Understanding clonal architecture provides important insights into tumor evolutionary dynamics and the timing of key genomic events during TGCT development.

Compared to seminomas, non-seminomas exhibited a significantly higher proportion of tumors with detectable subclonal populations (89% vs. 63%; Fisher’s exact test, *P* = 2.29e-06; OR = 4.7; **Fig. 5a**). This difference remained robust even when restricting the analysis to tumors with sufficient power for subclone detection^65^, as defined by the number of reads per chromosome copy (NRPCC) above 10 (*P*=8.15e-06; OR=9.9; **Supplementary Fig. 16**; **Supplementary Data 7**). Consistently, non-seminomas harbored a greater fraction of subclonal mutations (linear regression adjusted for tumor purity, *P*=0.0034; **Fig. 5b**), whereas seminomas were characterized by a predominance of clonal mutations (*P*=0.017). These observations reflect the higher genomic complexity and heterogeneity in non-seminomas relative to seminomas. Notably, the distribution of clonal versus subclonal mutations, as well as early versus late clonal mutations, was largely consistent across different mutational signatures (**Supplementary Fig. 17**), suggesting that diverse mutational processes remained active across both early and late stages of TGCT evolution.

**Fig. 5:**
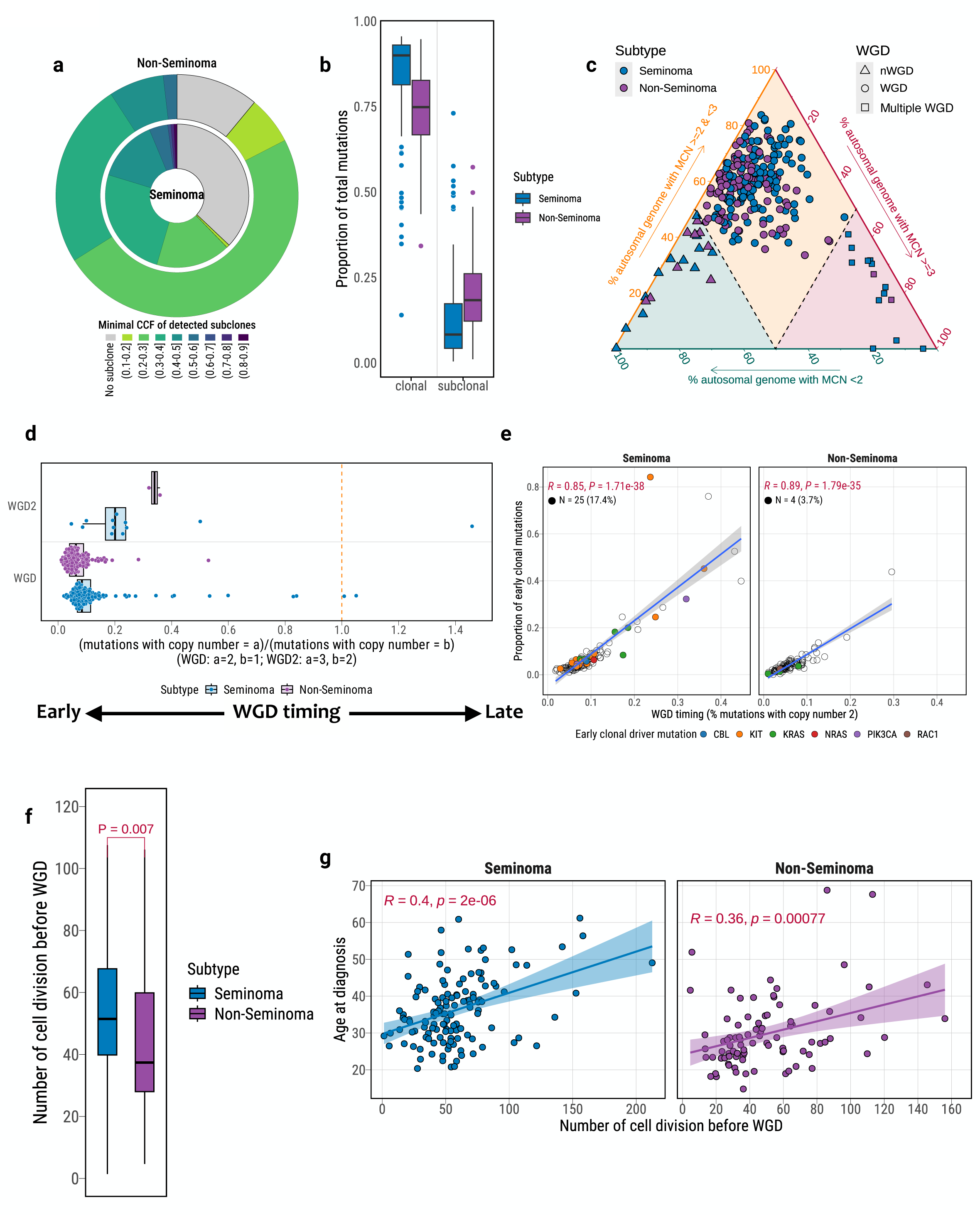
Early and Widespread Whole-Genome Doubling (WGD) and Clonal Architecture in TGCTs. (**a**) Distribution of tumors with minimal cancer cell fraction (CCF) of subclones in seminomas and non-seminomas, with non-seminomas showing higher intratumoral heterogeneity. (**b**) Comparison of clonal versus subclonal mutation proportions between seminomas and non-seminomas. (**c**) Ternary plot illustrating genome-wide major copy number (MCN) composition to infer WGD status. Each point represents a tumor, colored by subtype and shaped by WGD category. Tumors are classified as non-WGD, single WGD, or multiple WGD based on the relative proportions of the autosomal genome with MCN = 1, 2, or ≥3. (**d**) Timing of WGD events relative to the accumulation of clock-like mutations (SBS1 and SBS5). Boxplots show the ratio of mutations in genomic segments with major copy number (MCN) = 2 versus MCN = 1 (indicative of single WGD), or MCN = 3 versus MCN = 2 (indicative of multiple WGDs). High mutation ratios in duplicated segments suggest that WGD occurred early during tumor evolution in both seminomas and non-seminomas. (**e**) Correlation between WGD timing and early clonal mutation proportion in seminomas (left) and non-seminomas (right). Tumors harboring early clonal driver mutations prior to WGD are annotated by color, with counts and frequencies indicated above the plots. Pearson R and P-value are shown above scatterplots. (**f**) Estimated cell divisions before WGD in seminomas versus non-seminomas, inferred from pre-WGD mutation burden. P-values from two-sided Wilcoxon rank-sum tests are shown above boxplots. (**g**) Correlation between pre-WGD cell divisions and patient age at diagnosis in seminomas (left) and non-seminomas (right). Pearson R and P-value are shown above scatterplots. All boxplots display median, IQR, and whiskers extending to 1.5× IQR.

### Early and Widespread Whole-Genome Doubling as a Foundational Event in TGCT Evolution

Whole-genome doubling (WGD), a hallmark of cancer evolution, has been previously reported as an early event in TGCT development^18–20,66^. To further understand its role in shaping the distinct clonal architecture of TGCT subtypes, we investigated the prevalence and evolutionary timing of WGD in seminomas and non-seminomas. Remarkably, WGD was pervasive across 252 TGCTs, detected in 91.7% of tumors (91.6% of seminomas and 91.7% of non-seminomas; **Fig. 5c**; **Supplementary Data 8**), ranking TGCTs among the most WGD-enriched cancer types in TCGA^67^. Additionally, evidence of multiple WGD events was found in 6.35% of TGCT tumors, predominantly in seminomas (85.7%; *P*=0.027; OR=4.9), consistent with prior cytogenetic studies^68^ highlighting polyploidization as a hallmark of seminoma development.

Timing analysis revealed that WGD consistently occurred as a very early event in both TGCT subtypes, preceding the majority of somatic alterations (**Fig. 5d**). This observation aligns with previous reports that tetraploidization represents a defining early event in TGCT tumorigenesis^19,20^. Interestingly, we also observed a strong correlation between the relative timing of WGD and the proportion of early clonal mutations in both subtypes, supporting a key role for WGD in shaping subsequent tumor evolution (**Fig. 5e**). While WGD was typically among the earliest events in TGCTs, a subset of seminomas (N=25, 17.4%) harbored driver mutations in *KIT*, *KRAS*, *NRAS*, or *PIK3CA* prior to WGD, irrespective of WGD timing. In contrast, only three non-seminoma tumors showed evidence of pre-WGD driver mutations (three *KIT* and one *RAC1*). We further examined the temporal relationship between WGD and high-level copy number gains of chromosome 12p, a defining feature of TGCTs. Among tumors with sufficient resolution (N=33; **Supplementary Fig. 18-19**), chromosome 12p amplification was inferred to occur prior to WGD in 36% of cases, with enrichment in non-seminomas (*OR*=1.8; *P*=0.7), although this difference was not statistically significant. These results suggest that, although WGD generally marks the onset of tumorigenesis in TGCTs, driver mutations or 12q gain may occasionally precede WGD.

To estimate the developmental timing of WGD, we inferred the number of cell divisions prior to genome doubling based on pre-doubling mutation burdens and established mutation rates within primordial germ cells (PGCs)^19,20,69^. Across TGCTs, WGD was estimated to occur after approximately 49 cell divisions, supporting its likely origin during fetal germ cell development and consistent with its hypothesized role in the transformation of primordial germ cells into pre-invasive germ cell neoplasia *in situ* (GCNIS)^5,18^. Notably, non-seminomas exhibited significantly fewer pre-WGD cell divisions than seminomas (median: 39 vs. 54; *P*=0.007; **Fig. 5f**), suggesting that WGD occurred at an earlier developmental stage in non-seminomas. Moreover, the number of pre-WGD cell divisions was significantly correlated with patient age at diagnosis in both subtypes (**Fig. 5g**), indicating that the developmental timing of WGD may influence tumor initiation and clinical presentation.

These findings further support WGD as an early event in TGCT tumorigenesis, providing a shared genomic foundation for both seminomas and non-seminomas. However, the timing of WGD and the occurrence of pre-WGD driver mutations and 12q gain differ between subtypes, contributing to their distinct evolutionary trajectories and biological behaviors. These insights highlight WGD not only as a hallmark of TGCT tumorigenesis but also as a critical determinant of subsequent genomic evolution, heterogeneity, and disease onset.

### Distinctive and Germ Cell-Like Transcriptomic Program Uniquely Defines TGCTs Among Human Cancers

We next investigated whether these tumors also exhibit a uniquely defined transcriptomic landscape compared to other solid tumors. To this end, we performed single-sample Gene Set Enrichment Analysis (ssGSEA) across all TCGA cancer types using RNA-seq data, focusing on both Hallmark and KEGG gene sets (**Supplementary Data 9-10)**. Strikingly, TGCTs showed the highest enrichment scores across multiple pathways related to cell cycle regulation (*MYC_TARGETS*, *E2F_TARGETS*, *MITOTIC_SPINDLE*), spermatogenesis, and DNA repair (*DNA_REPAIR*, *HOMOLOGOUS_RECOMBINATION*, *BASE_EXCISION_REPAIR*, *MISMATCH_REPAIR*) (**Fig. 6a-d**). In contrast, TGCTs exhibited notably low activity in multiple metabolism-related pathways and TNFA_SIGNALING_VIA_NFKB, suggesting minimal inflammatory and metabolic signaling compared to other cancer types.

**Fig. 6:**
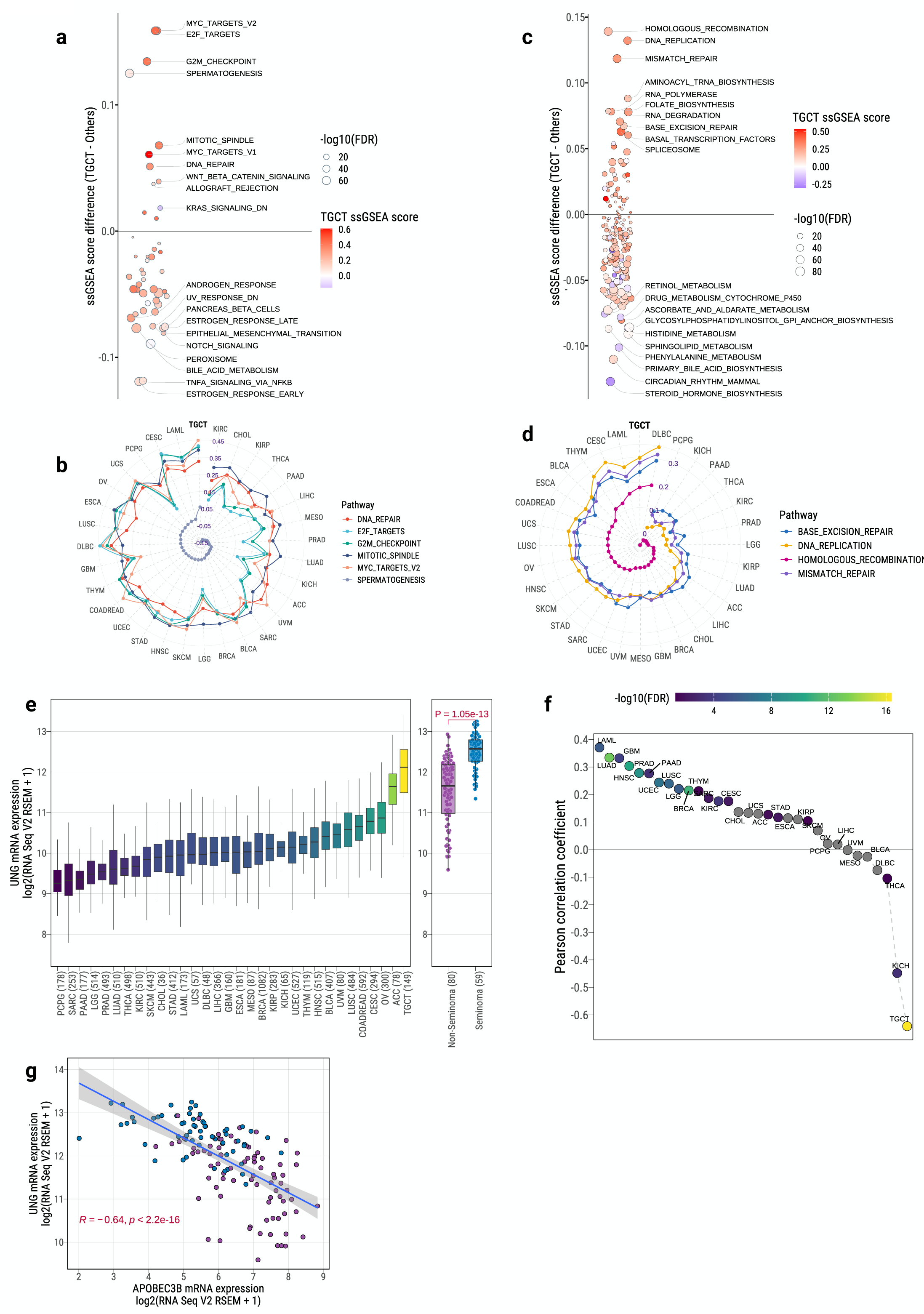
Germ Cell-Like Transcriptomic Program and Unique APOBEC Mutagenesis Suppression in TGCTs. (**a**) Differential enrichment of Hallmark gene sets in TGCTs versus other TCGA tumor types using ssGSEA scores. Enriched pathways include cell cycle, DNA repair, and spermatogenesis; depleted pathways include metabolic and immune-related processes. Circle size indicates−log10 FDR from two-sided Wilcoxon rank-sum tests; color denotes median ssGSEA score enrichment in TGCTs. Benjamini–Hochberg correction was applied. (**b**) Radar plot of TGCT-specific enrichment for select Hallmark pathways (SPERMATOGENESIS, DNA_REPAIR, E2F_TARGETS, G2M_CHECKPOINT, MITOTIC_SPINDLE, MYC_TARGETS_V2) across TCGA tumor types. (**c**) Differential enrichment of KEGG gene sets in TGCTs, with top pathways including HOMOLOGOUS_RECOMBINATION, MISMATCH_REPAIR, and BASE_EXCISION_REPAIR. Circle size indicates −log10 FDR from two-sided Wilcoxon rank-sum tests; color denotes median ssGSEA score enrichment in TGCTs. Benjamini–Hochberg correction was applied. (**d**) Radar plot of ssGSEA enrichment for core DNA repair KEGG pathways across TCGA tumor types, with TGCTs ranking highest. (**e**) *UNG* (uracil DNA glycosylase) mRNA expression across TCGA tumors, with TGCTs showing the highest levels and seminomas higher than non-seminomas. P-values from two-sided Wilcoxon rank-sum tests are shown above boxplots. *Boxplots display median, IQR, and whiskers extending to 1.5× IQR*. (**f**) Pearson correlation between *APOBEC3B* and *UNG* expression across TCGA tumors, with TGCTs showing a unique negative correlation. FDR from Benjamini–Hochberg correction is shown. (**g**) Scatterplot of negative association between *APOBEC3B* and *UNG* mRNA expression in TGCTs. Pearson R and P-value are shown below.

Among DNA repair genes, *UNG* (uracil DNA glycosylase), a key enzyme initiating base excision repair (BER)^70^, showed the highest expression in TGCTs among all TCGA tumors (**Fig. 6e**). Moreover, seminomas displayed significantly higher *UNG* expression than non-seminomas (Wilcoxon test, *P*=1.05e-03; fold change=5.3), highlighting enhanced BER activity particularly in this subtype. Interestingly, despite widespread APOBEC mutagenesis observed across many cancer types, both seminoma and non-seminoma uniquely lacked any APOBEC-related mutational signatures, as confirmed by both mutational signature analysis and motif-based evaluations. While most tumors showed a strong positive correlation between the APOBEC mutator APOBEC3B and UNG expression, TGCTs exhibited a significant negative correlation (*FDR*=4.34e-17; Pearson *R*=-0.64; **Fig. 6f-g**). This unique pattern may reflect an early developmental transcriptional environment, possibly linked to the primordial germ cell origin of TGCTs^71^, that does not support APOBEC3B activity and mutagenesis.

We further explored transcriptional differences between TGCT subtypes. Non-seminomas were significantly enriched for pathways associated with ANGIOGENESIS, EPITHELIAL_MESENCHYMAL_TRANSITION (EMT), NOTCH_SIGNALING, and HYPOXIA, consistent with their more aggressive and differentiated phenotype. In contrast, seminomas exhibited higher activity in INTERFERON_RESPONSE, SPERMATOGENESIS, and KRAS_SIGNALING pathways (**Supplementary Fig. 20**), reflecting their undifferentiated, germ-cell-like state.

Together, these findings reveal that TGCTs harbor an exceptionally distinctive transcriptomic program, characterized by high cell cycle activity, strong germ cell features, elevated DNA repair processes, but minimal metabolic and immune signaling. These transcriptomic patterns align closely with pathways implicated by TGCT susceptibility loci identified in genome-wide association studies^29,30^.

This profile sharply distinguishes TGCTs from all other solid tumors within the TCGA cohort and underscores their unique developmental origin and biology. Moreover, this germ cell-like transcriptional landscape may represent a key vulnerability and inform TGCT-specific therapeutic strategies.

## Discussion

Our integrative genomic characterization of TGCTs sheds light on the fundamental molecular processes underlying the distinct evolutionary trajectories of seminomas and non-seminomas, extending beyond previous targeted and exome sequencing studies. In addition to confirming canonical drivers and the hallmark amplification of chromosome 12p, we identified several infrequently mutated but potentially relevant driver genes—such as *FAT4*, *GNAS*, *LRP1B*, *RPL5*, and *TET1*—many of which are involved in chromatin remodeling, growth signaling, or translational regulation. While previous TCGA studies using SNP arrays highlighted broad SCNA patterns in TGCTs^5^, deep WGS allowed us to refine the resolution of these alterations and uncover novel, high-confidence focal events, even on sex chromosomes. Notably, we identified a top significant and previously underappreciated focal deletion at Xq28 encompassing *SPRY3*, a potential negative regulator of RAS/MAPK signaling^72^. As *SPRY3* resides within the PAR2 on both the X and Y chromosomes, its dosage and regulation are typically shared between sex chromosomes. However, the Y-linked allele is often transcriptionally silent^73^, and our observation of frequent X-linked *SPRY3* deletions in TGCTs suggests a loss of function that could exacerbate MAPK pathway activation. We also observed recurrent focal deletions of *RBMY* family genes on the Y chromosome, which encode RNA binding proteins critical for spermatogenesis, occurring predominantly in seminomas. This pattern suggests that instability of the sex chromosomes may disrupt male germline maintenance and highlights distinct lineage-specific vulnerabilities between TGCT subtypes.

Chromosome X amplification and subsequent inactivation via XIST represent another major genomic hallmark of TGCTs. We observed subtype-specific strategies in managing X chromosome dosage: seminomas maintain constitutive *XIST* expression and epigenetic silencing, leading to increased replication stress, whereas non-seminomas modulate *XIST* based on chromosome copy number and show greater epigenetic plasticity, reflecting their more differentiated state. This suggests a potential vulnerability in seminomas related to replication stress, while non-seminomas may possess adaptive mechanisms mitigating such genomic instability. In addition, our data reinforce the near-universal prevalence of whole-genome doubling (WGD) as a foundational event in TGCT development, observed in over 90% of tumors. Our clonal timing analyses indicate that WGD occurs exceptionally early—preceding most other somatic alterations—supporting a model in which WGD represents the first major genomic "hit" during the earliest stages of tumorigenesis. This is consistent with the fetal origin of TGCTs, where the initial transformation of primordial germ cells (PGCs) into germ cell neoplasia *in situ* (GCNIS) is thought to occur *in utero*. Based on our data and consistent with prior studies^5,18,74^, WGD likely arises at this prepubertal GCNIS stage. In contrast, events such as 12p gain, and acquisition of additional driver mutations appear to occur later, potentially during the GCNIS-to-invasive tumor transition that aligns with postpubertal germ cell reactivation. Notably, GCNIS lesions typically lack 12p amplification^18,75^, further supporting this temporal distinction. Moreover, our finding that WGD timing correlates with patient age at diagnosis suggests that differences in the timing or dynamics of PGC expansion and dormancy may shape tumor onset. Interestingly, a subset of seminomas (17.4%) harbored activating driver mutations that occurred prior to WGD. Consistent with a previous report^5^, we observed a small number of early *KIT* mutations. However, with the larger sample size we were additionally able to identify a small number of *RAS* mutations that appear to precede WGD. These early events may represent an alternative or complementary route to tumor initiation in some seminomas, potentially priming GCNIS cells for transformation prior to genome duplication. Together, these findings offer a refined developmental framework for TGCT pathogenesis, linking early mutational and genomic events to in utero origins, and supporting a model in which both timing and sequence of alterations shape distinct evolutionary trajectories across subtypes.

While seminomas harbored more frequent driver mutations and focal amplifications, non-seminomas exhibited significantly greater subclonal diversity and structural rearrangements, underscoring their increased intratumoral heterogeneity. This dichotomy highlights divergent evolutionary trajectories, with seminomas maintaining a more germ cell–like and genomically stable profile, while non-seminomas transitioning toward greater genomic complexity and heterogeneity. Mutational signatures analysis further delineated this divergence. Most notably, we identified SBS87, previously reported only in relapsed acute lymphoblastic leukemia and experimentally linked to thiopurine exposure^51,76^, as highly prevalent in non-seminomas. This signature is completely absent from several large-scale WGS-based cancer genomics datasets, including PCAWG (4,645 whole genomes and 19,184 exomes)^77,78^, and the St. Jude pediatric cancer cohort (785 whole genomes)^79^. In a recent pan-cancer analysis of 12,222 whole-genome sequenced tumors^80^, SBS87 was detected in only a single breast cancer case. The unexpected presence of SBS87 in 86.2% of treatment-naive non-seminomas and nearly half of all TGCTs raises two plausible, but not mutually exclusive explanations: (1) prior environmental or pharmacologic thiopurine exposure, or (2) an endogenous mutagenic process that phenocopies thiopurine-induced DNA damage. Such a process may reflect germ-cell-specific aspects of DNA damage and repair, and could also potentially be influenced by *in utero* factors, including maternal exposures during germ cell development. Thiopurines, including azathioprine and mercaptopurine, are widely used as immunosuppressive agents in the treatment of inflammatory bowel disease (IBD) and other autoimmune disorders. Experimental studies have demonstrated their detrimental effects on testicular function and male fertility^55^, although clinical evidence remains limited^54^. IBD itself has been linked to an elevated risk of intestinal tract cancers^81–83^, and thiopurine usage has been associated with increased risk of multiple cancers^84^. Although isolated case reports have described TGCT development following thiopurine treatment^85,86^, there is currently no evidence supporting an association between thiopurine exposure and TGCT risk. Although population-scale datasets such as the UK Biobank and AllofUs included a small number of individuals with both TGCT and IBD or documented thiopurine use (fewer than 10 cases), we found no evidence of an association between thiopurine exposure and TGCT risk; Given the prevalence of SBS87 in treatment-naive tumors, a purely treatment-related origin seems unlikely. Instead, we propose that germ cell tumors may harbor intrinsic vulnerabilities in purine metabolism or DNA repair that generate thiopurine-like lesions. These vulnerabilities may be further influenced by maternal exposures during early germ cell development, potentially contributing to an endogenous-like mutational process that mimics thiopurine-induced damage. The germline context of TGCTs, characterized by global epigenetic reprogramming, replication stress, and altered oxidative metabolism, may also amplify susceptibility to such damage. Thus, SBS87 may reflect a germ cell-specific mutational footprint shaped by developmental and cellular context, rather than direct thiopurine exposure. Indeed, SBS87 showed evolutionary dynamics similar to clock-like signatures SBS1 and SBS5 (**Supplementary Fig. 17**), suggesting SBS87 mutations may accumulate steadily over time. This pattern is perhaps more consistent with the hypothesis of an endogenous origin. However, given the low overall mutation burden in TGCTs, clonal timing estimates carry uncertainty and should be interpreted cautiously. Further mechanistic and experimental studies will be essential to clarify the origin and implications of SBS87 in germ cell tumorigenesis. Importantly, SBS87 is associated with extended tumor latency and telomere elongation in non-seminomas, suggesting it influences not only tumor initiation but also progression and clinical phenotype. These findings underscore the need to further explore the endogenous and environmental determinants of germ cell mutagenesis, and their long-term implications in testicular tumorigenesis.

Clinically, these insights could inform strategies for personalized therapy, especially for recurrent or refractory disease. Targeting replication stress vulnerabilities in seminomas, or exploiting telomere maintenance mechanisms in SBS87-enriched non-seminomas, represents promising avenues for therapeutic intervention.

In conclusion, our comprehensive analysis significantly advances our understanding of TGCT biology, highlighting early genome-wide events, distinct subtype evolutionary pathways, and unique molecular vulnerabilities. These findings not only refine the molecular classification of TGCT subtypes but also provide actionable insights for novel therapeutic approaches and biomarker development.

## Methods

### Whole-genome sequencing (WGS) data processing

We analyzed WGS data from 252 primary testicular germ cell tumors with matched blood samples, comprising 143 seminomas and 109 non-seminomas, collected as part of The Cancer Genome Atlas (TCGA) project. Aligned sequencing data in BAM format were downloaded from the Genomic Data Commons (GDC) Data Portal (https://portal.gdc.cancer.gov/). All samples were sequenced using 2 x 151 bp paired-end reads. Corresponding clinical and exposure metadata were also retrieved from the GDC Data Portal for integrative analyses. To preserve read group, sequencing lane metadata, and unmapped reads, BAM files were converted to FASTQ format using Bazam (v1.0.1)^87^. Reads were aligned to the GRCh38 human reference genome using the BWA-MEM algorithm. The resulting alignments were processed into analysis-ready CRAM files using the previously described Sherlock-*Lung* pipeline ^58,88–90^. This included marking of duplicate reads, local realignment around indels, and base quality score recalibration. Quality control was performed on the resulting CRAM files. Sequencing metrics were collected using Picard (v2.23.3), depth of sequencing was estimated using mosdepth^91^ (v0.3.3), tumor-normal sample concordance was assessed using Somilar (v0.2.6), and genetic ancestry was evaluated using VerifyBamID^92^ (v2.0.1). One tumor sample failed QC and was excluded from downstream analyses.

### Analysis of Genome-Wide Somatic Alterations

To identify genome-wide somatic alterations in TGCTs, we applied the same high-confidence pipelines used in our Sherlock-*Lung* study^58,88–90^, leveraging the comparable sequencing depth of TCGA-TGCT WGS data. Somatic single nucleotide variants (SNVs) and small insertions/deletions (indels) were called using an ensemble of four variant callers: Strelka (v2.9.10), MuTect, MuTect2, and TNscope (Sentieon Genomics v2023.08.03). A consensus mutation set was generated by retaining only variants called by at least three of the four algorithms^58^. To filter potential germline contamination, variants were excluded if present at allele frequency >0.001 in the Genome Aggregation Database (gnomAD v4.0). Functional annotation of somatic variants was performed using ANNOVAR (v2020-06-08). To identify candidate cancer driver genes, we employed the IntOGen pipeline^21^ (v2020.02.0123), which integrates seven orthogonal algorithms to detect signals of positive selection based on mutational clustering, recurrence, and functional impact.

Allele-specific somatic copy number alterations (SCNAs) were inferred using the NGSpurity pipeline^93^, which includes Battenberg^94^ (v2.2.9), DPClust^94,95^ (v2.2.8), and ccube^96^ (v1.0) algorithms. Tumor purity, ploidy, and clonal architecture were jointly estimated. SCNA profiles failing manual quality inspection were iteratively re-estimated by adjusting purity and ploidy, guided by local copy number status and validated by manual review. GISTIC2.0 was used to identify recurrent focal SCNAs using the major clonal copy number state for each segment.

Structural variants (SVs) were identified using Meerkat^97^ (v0.189) and Manta^98^ (v1.6.0), and a union set of SVs passing tool-recommended filters was used for downstream analysis^59^. Gene fusions were filtered based on Meerkat annotations with features including ’gene-gene’, ’head-tail’, and ’in-frame’ or ’out-of-frame’ fusion orientation.

To detect somatic insertions of transposable elements (TEs), we applied TraFiC-mem^99^ (v1.1.0), retaining TE insertions that passed default quality and filtering criteria.

### Copy Number Analysis of Chromosome X and Y

SCNA on chromosome X were analyzed using a segmentation-based approach adapted from the Battenberg algorithm^94^, with specific modifications to account for the hemizygous nature of the non-pseudoautosomal (non-PAR) region in males. GC-corrected LogR values were segmented using piecewise constant fitting, incorporating structural variant (SV) breakpoints where available to improve segmentation accuracy. LogR values were calibrated using autosomal diploid and gained regions to mitigate baseline noise. Segment-level copy number states were then inferred by comparing observed LogR values to a model of expected signal intensities across a range of integer copy number states, adjusted for tumor-specific purity and ploidy estimates. Clonality was determined by assessing deviations from expected LogR values, enabling classification of each SCNA as clonal or subclonal. For subclonal segments, cancer cell fraction (CCF) was estimated based on allele-specific signal patterns. This approach allowed high-resolution, allele-aware mapping of chromosome X SCNAs, including detection of focal gains, losses, and clonality status across the non-PAR region.

Copy number analysis of chromosome Y posed technical challenges due to the extensive heterochromatic region (Yq12) and the lack of support for Y chromosome segmentation in the Battenberg algorithm. Therefore, we employed CNVkit^100^ to estimate copy number states on chromosome Y, adjusting for refined tumor purity and using the haploid male genome as a reference. This approach enabled reliable detection of large-scale gains and losses across the euchromatic regions of chromosome Y in TGCT samples.

### Inferring Copy Number Gain of Chromosome X via Sequencing Depth

To assess copy number gain of chromosome X, we estimated the mean sequencing depth for chromosome X and the autosomes in each sample. Depth was calculated as the total number of mapped reads multiplied by read length, divided by the total mappable length of the chromosome. The ratio of chromosome X to autosomal sequencing depth was used to infer relative gains or losses of chromosome X. In normal male blood samples, this ratio was centered around 0.5, consistent with a single copy of chromosome X in male. In tumor samples, this ratio was adjusted for tumor purity to account for normal cell contamination. The adjusted X-to-autosome depth ratio *R* was calculated as:

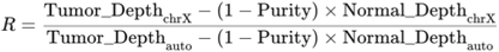

Where: Tumor_Depth_chrX_: observed mean sequencing depth of chromosome X in tumor. Normal_Depth_chrX_: mean depth of chromosome X in matched normal sample. Tumor_Depth_auto_: mean sequencing depth of autosomes in a tumor. Normal_Depth_auto_: mean autosomal depth in normal. Purity: estimated tumor purity from allele-specific copy number analysis.

This approach enabled robust estimation of relative chromosome X dosage across TGCT tumors. A similar method was applied to evaluate potential gain or loss of chromosome Y.

### Mutational signature analysis

Somatic mutation matrices were generated using SigProfilerMatrixGenerator^101^ to classify single base substitutions (SBS), indels (ID), and doublet base substitutions (DBS) across all samples. Due to the low burden of DBS mutations in TGCTs (median = 5 DBS per tumor), mutational signature analysis was limited to SBS and ID classes. De novo extraction of SBS and ID signatures was performed using SigProfilerExtractor^47^ (v1.1.21), applying default parameters and using SBS-96 and ID-83 mutation contexts, respectively. Extracted signatures were subsequently decomposed into known COSMIC v3.4 reference signatures^102^ based on the GRCh38 genome build.

To estimate mutational signature activities at the individual sample level, we applied the Mutational Signature Attribution (MSA) tool^48^, which uses a non-negative least squares (NNLS) algorithm optimized via configurable simulations. MSA was used to fit the COSMIC signature set (as determined by SigProfilerExtractor) to each tumor, with attribution thresholds automatically optimized by the MSA pipeline. A parametric bootstrap procedure was employed to generate 95% confidence intervals for each signature’s activity.

### Timing mutations and copy number gains using MutationTimeR

To infer the temporal ordering of somatic events, we used the R package MutationTimeR^103^ (v1.00.2) to estimate the timing of mutations relative to clonal and subclonal copy number states, as well as to compute the relative timing of copy number gains^103^. Input data included high-confidence somatic single-nucleotide variants (SNVs) and indels, along with final allele-specific copy number profiles derived from SCNA analysis.

MutationTimeR classifies mutations into four categories based on their timing relative to copy number alterations: early clonal (occurring before a clonal gain), late clonal (after a clonal gain), clonal N/A (ambiguous timing), and subclonal (arising in a subclonal population). Early clonal mutations were used to infer events that occurred prior to copy number gains, enabling reconstruction of the temporal sequence of tumor evolution.

### Chronological Timing Analysis and Estimation of Tumor Latency

To infer the timing of tumor evolution and estimate latency, we applied a chronological time analysis framework established in prior pan-cancer studies^103,104^. This approach enables estimation of the age at which the most recent common ancestor (MRCA) of the tumor emerged, as well as the latency period between MRCA emergence and clinical diagnosis.

We implemented a hierarchical Bayesian linear regression model to relate the number of clock-like CpG>TpG mutations (at NpCpG sites) to patient age at diagnosis. This model adjusts for tumor ploidy, purity, and subclonal composition, following the methodology developed by the PCAWG Evolution and Timing Working Group^103^ (https://github.com/gerstung-lab/PCAWG-11/). The model infers the MRCA emergence time by leveraging the linear accumulation of SBS1 mutations, using a tumor acceleration rate of 1x, appropriate for TGCTs with low mutational burden compared to other cancer types. Tumor latency was defined as the time interval between MRCA emergence and age at diagnosis, calculated by subtracting the estimated MRCA age from the observed age at diagnosis for each patient.

### Identification and Timing of Whole-Genome Doubling

To determine whole-genome doubling (WGD) status, we applied criteria established by Bielski et al67. Tumors were classified as WGD-positive if >50% of the autosomal genome exhibited a major copy number (MCN)—defined as the copy number of the more abundant allele in each segment—of ≥2. Additionally, at least 11 autosomes were required to contain MCN□≥□2 across >50% of their segments. For identification of tumors with evidence of multiple WGD events, we applied a more stringent threshold, requiring >50% of the autosomal genome to have MCN□≥□3.

To assess the evolutionary timing of WGD, we calculated the ratio of somatic mutations occurring in segments with MCN = 2 versus MCN = 1 (indicative of single WGD), or MCN = 3 versus MCN = 2 (indicative of multiple WGD events). Higher mutation ratios in duplicated segments are interpreted as evidence of early WGD. For this analysis, we restricted the mutation set to clock-like substitutions (SBS1 and SBS5), which are known to accumulate at a relatively constant rate throughout tumor evolution.

The number of cell divisions preceding WGD was estimated using a previously published molecular clock model^69^, which assumes a mutation rate of 0.5–0.7 substitutions per haploid genome per cell division within primordial germ cells (PGCs). This approach assumes that the majority of detectable somatic mutations arise post-PGC specification, and that early embryonic mutations are not captured in bulk sequencing data^20^.

To further resolve the temporal order of WGD relative to recurrent focal amplifications—such as those involving chromosome 12p—we applied the AmplificationTimeR algorithm^105^ (v1.1.2). This method infers the relative timing of copy number gains based on the observed mutation multiplicity and the segment’s highest copy number. As with WGD timing, we limited this analysis to SBS1 and SBS5 mutations to preserve clock-like behavior in the model’s assumptions.

### Isochromosome 12p Classification

Canonical isochromosome 12p [i(12p)] events were identified using allele-specific somatic copy number profiles. A tumor was classified as harboring i(12p) if it exhibited at least one extra copy of chromosome arm 12p relative to 12q, consistent with gain through a centromeric duplication mechanism. When p and q arms were segmented separately, we required that the largest 12p segment exceeded the largest 12q segment by at least one copy.

To assess the robustness of this classification, we applied a more stringent threshold, requiring at least two additional copies of 12p relative to 12q. Under this criterion, 94% of tumors previously classified as i(12p)-positive retained their designation, confirming the stability of our detection strategy.

### Detection of Extrachromosomal DNA (ecDNA)

Candidate extrachromosomal DNA (ecDNA) structures were identified from tumor BAM files using the AmpliconSuite-pipeline^106^ (v1.3.7), which performs all necessary preprocessing steps, including read alignment, copy number variation (CNV) calling, and seed interval detection, prior to ecDNA reconstruction. Amplicon structures were then resolved using AmpliconArchitect^31^. Final classification of amplicons was performed using AmpliconClassifier^106^, which distinguishes ecDNA from other amplification mechanisms such as breakage–fusion–bridge (BFB) cycles, linear amplifications, and complex rearrangements.

### RNA-Seq and DNA Methylation Data Analysis

PanCancer RNA-Seq data for TCGA samples were obtained from cBioPortal (study ID: *TCGA, PanCancer Atlas*), using the "mrna_seq_v2_rsem" dataset for gene expression quantification. For comparisons between TCGA testicular germ cell tumors and normal testis tissue, we retrieved preprocessed data from the UCSC Xena Browser (dataset ID: *TCGA-GTEx-TARGET-gene-exp-counts.deseq2-normalized.log2*). For cross-cancer comparisons of *XIST* expression, only male TCGA subjects were included in the analysis.

Replication stress scores were computed based on a weighted principal component–derived signature from a previously published study^46^. Seventeen replication stress–associated genes were included in the score. For each sample, gene expression values (Z score–normalized across all genes within a sample set) were multiplied by corresponding gene weights and summed to yield a replication stress score. Final scores were Z-score normalized across all samples within each analysis cohort.

To characterize the transcriptomic landscape of TGCTs relative to other cancer types, we performed single-sample Gene Set Enrichment Analysis (ssGSEA) across all TCGA tumor types using the GSVA R package^107^. Gene sets were derived from the *Molecular Signatures Database (MSigDB)* via the msigdbr R package, including both Hallmark and KEGG collections. ssGSEA scores were compared between TGCTs and other TCGA cancer types, as well as between TGCT subtypes, using two-sided Wilcoxon rank-sum tests.

For DNA methylation analyses, preprocessed Level 3 beta values for the TCGA TGCT cohort were downloaded from the GDC Data Portal. To compare methylation across chromosomes, we calculated the median beta value for CpG probes located on chromosome X and autosomes, stratified by TGCT subtype. Given the globally hypomethylated nature of TGCTs, we restricted the analysis to informative loci by including only CpG probes with a median absolute deviation (MAD) > 0.1 across all samples.

### Statistical analyses and data visualization

All statistical analyses were conducted using R software (v4.3.2) (https://www.r-project.org/). Data visualizations were generated using the *ggplot2* package (https://ggplot2.tidyverse.org/) along with other supporting R packages. For comparisons between two groups of continuous variables, the two-sided Mann–Whitney U test (Wilcoxon rank-sum test) was used. The two-sided Fisher’s exact test was applied for enrichment analyses involving categorical variables. Where multiple testing was performed, false discovery rate (FDR) correction was applied using the Benjamini–Hochberg method. A p-value or FDR < 0.05 was considered statistically significant. Linear and logistic regression models were used to assess associations between genomic features or alterations and TGCT subtypes, adjusting for tumor purity and age when appropriate.

## Data availability

Access to TCGA controlled data can be requested through dbGaP (study accession: phs000178.v11.p8). Raw whole-genome sequencing (WGS) data (BAM files) for the TCGA TGCT cohort were obtained from the Genomic Data Commons (GDC) data portal (https://portal.gdc.cancer.gov/) under standard TCGA data access policies. Processed RNA-seq data (quantified as RSEM values) for the TCGA TGCT cohort are available via cBioPortal (https://www.cbioportal.org/), using the study name "Testicular Germ Cell Tumors (TCGA, PanCancer Atlas)." Integrated RNA-seq data combining TCGA TGCT and GTEx testicular tissues can be accessed through the Xena browser (https://xenabrowser.net/) using dataset ID: TCGA-GTEx-TARGET-gene-exp-counts.deseq2-normalized.log2. Level 3 DNA methylation data (beta values) for the TCGA TGCT cohort are also accessible from the GDC data portal.

## Code availability

The bioinformatics pipelines used for whole-genome sequencing analysis are available at https://github.com/xtmgah/Sherlock-Lung. The source code used to generate the main figures presented in this manuscript is accessible at https://github.com/xtmgah/TCGA-WGS-Mansucripts.

## Acknowledgments

This work was supported by the Intramural Research Program of the National Cancer Institute, US National Institute of Health (NIH). This work utilized the computational resources of the NIH HPC Biowulf cluster (https://hpc.nih.gov).

## Author Contributions

Conceptualization, KB, KLN, TZ; Methodology, TZ; Formal Analysis, TZ, JOW, KB, JAW; Resources, KB, SJC, TZ; Data Curation, TZ, SJC; Writing – Original Draft, KB, TZ; Writing – Review & Editing, KB, JZ, AMM, MZ JOW, JAW, HZ, CL, WL, BZ, SJC, KLN, TZ; Visualization, TZ; Supervision, TZ.

## Inclusion & Ethics Statement

This study utilized publicly available genomic, transcriptomic, and epigenomic data from The Cancer Genome Atlas (TCGA), ensuring broad transparency and equitable access to resources. No new human subjects or biological samples were collected. All analyses were performed in compliance with the TCGA data usage policies and NIH ethical guidelines.

We acknowledge the contributions of the diverse TCGA participant cohort and the researchers who generated and curated these datasets. Authorship reflects contributions across scientific roles including data analysis, interpretation, and manuscript preparation.

## Supplementary Figures

**Supplementary Fig. 1:** Summary of clinical and sequencing characteristics for the TCGA TGCT cohort. (**a**) Comparison of age at diagnosis between non-seminomas and seminomas. P-value from two-sided Wilcoxon rank-sum test is shown. (**b**) Distribution of median WGS sequencing depth for tumor and matched normal samples. Median depth across samples of each type is indicated by a red vertical line. (**c**) Principal component analysis (PCA) for ancestry inference using 1000 Genomes reference samples.

**Supplementary Fig. 2:** Stacked bar plot showing the timing of point mutations in recurrent driver genes in TGCTs. The majority of mutations occur early during tumor evolution in both seminomas and non-seminomas.

**Supplementary Fig. 3:** Lollipop plots of non-synonymous mutations in the major TGCT driver genes *KIT* (**a**) and *KRAS* (**b**). Mutation counts are shown separately for seminomas (top) and non-seminomas (bottom), generated using ProteinPaint (https://proteinpaint.stjude.org/).

**Supplementary Fig. 4:** Arm-level SCNA classification of TGCTs. Left, Unsupervised clustering of TGCT samples based on arm-level SCNA profiles and visualization of copy number is based on relative copy number: total copy number – ploidy (non-WGD = 2 and WGD = 4). The analysis revealed three major SCNA subgroups, designated C1, C2, and C3. Top, Frequency of SCNA events, including gains (red), losses (blue), and copy-neutral loss of heterozygosity (LOH; black line) for each chromosomal arm.

**Supplementary Fig. 5:** GISTIC-based analysis of recurrent focal SCNAs for TCGT subtypes. (**a-b**) Significant focal amplifications in seminomas and non-seminomas, respectively. (**c**) Comparison of amplification significance (q-values) between subtypes. (**d-e**) Significant focal deletions in seminomas and non-seminomas, respectively. (**f**) Comparison of deletion significance between subtypes.

**Supplementary Fig. 6:** Detection and classification of extrachromosomal DNA (ecDNA) in TGCTs. (**a**) Comparison of median copy number among different amplification types (linear, ecDNA, BFB, and complex-non-cyclic). P-values from two-sided Wilcoxon rank-sum tests are shown above boxplots. *Boxplots display median, IQR, and whiskers extending to 1.5× IQR.* (**b**) Frequency of focal amplicons stratified by amplification mechanism. (**c–e**) Representative examples of ecDNA-driven focal amplifications involving *KIT* (**c**), *KRAS* (**d**), and *MDM2* (**e**), as reconstructed by the *AmpliconArchitect* algorithm. Edge colors indicate the orientation of junctions between genomic segments, and edge thickness reflects breakpoint copy number as estimated by *AmpliconArchitect*. Vertical dashed lines separate segments originating from different chromosomes, while dotted lines distinguish distinct regions within the same chromosome. Breakpoint edge numbers correspond to those used in the *AmpliconArchitect* reconstruction.

**Supplementary Fig. 7: Copy number analysis of chromosome Y.** (**a**) Tumor purity-adjusted sequencing depth ratios of chromosome Y vs. autosomes in matched normal blood, seminomas, and non-seminomas. P-values from two-sided Wilcoxon rank-sum tests. *Boxplots display median, IQR, and whiskers extending to 1.5× IQR.* b, (**b**) Copy number profiles of chromosome Y in TGCTs, separated by seminoma (left) and non-seminoma (right), clustered by total copy number. The y-axis indicates chromosome Y positions.

**Supplementary Fig. 8:** Distribution of DNA methylation levels (beta) on chromosome X versus autosomes, stratified by TGCT subtype.

**Supplementary Fig. 9:** Replication stress score comparisons. (**a**) Comparison of replication stress between seminomas and non-seminomas. (**b**) Association between replication stress and chromosome X copy number. P-values from linear regression adjusted for tumor purity. *Boxplots display median, IQR, and whiskers extending to 1.5× IQR*.

**Supplementary Fig. 10:** Differential expression analysis of X-linked genes between TGCT tumors with and without chromosome X amplification. Positive logD fold change indicates higher gene expression in tumors with chromosome X amplification. P-values were calculated using two-sided Wilcoxon rank-sum tests; FDR values were computed using the Benjamini–Hochberg method.

**Supplementary Fig. 11:** Correlation between patient age at diagnosis and number of clock-like mutations (SBS1 and SBS5), stratified by TGCT subtype. Pearson correlation coefficient (R) and P-values are shown.

**Supplementary Fig. 12:** Differential expression of thiopurine metabolism–related genes between seminomas and non-seminomas. (a) Volcano plot showing logD fold-change (x-axis) and statistical significance (–logDD FDR, y-axis) for 15 genes involved in thiopurine metabolism, comparing non-seminomas versus seminomas. Genes upregulated in non-seminomas are shown in purple, and those upregulated in seminomas are shown in blue. Dashed red line indicates FDR = 0.05. P-values from Wilcoxon rank-sum tests; FDR values calculated using Benjamini–Hochberg correction. (b) Boxplots showing normalized RNA-seq expression (logD[RSEM + 1]) for the same 15 genes across TGCT subtypes. All boxplots display median, interquartile range, and whiskers extending to 1.5× the IQR.

**Supplementary Fig. 13:** Association between tumor latency and major genomic alterations (>3% frequency). P-values from Wilcoxon rank-sum tests; FDR values calculated using Benjamini–Hochberg correction.

**Supplementary Fig. 14:** Correlation between SBS5 mutational burden and estimated tumor latency, stratified by TGCT subtype. Pearson R and P-values are shown.

**Supplementary Fig. 15:** Association between estimated tumor telomere length and major genomic alterations (>3% frequency). Linear regression adjusted for tumor purity and age at diagnosis. FDR values calculated using Benjamini–Hochberg correction.

**Supplementary Fig. 16:** Distribution of subclonal cancer cell fractions (CCFs) in tumors with sufficient sequencing power (NRPCC > 10), stratified by TGCT subtype.

**Supplementary Fig. 17:** Proportion of early clonal, clonal, late clonal, and subclonal mutations attributed to each SBS signature, stratified by TGCT subtype.

**Supplementary Fig. 18:** AmplificationTimeR timing analysis showing late WGD (W) relative to first gain of 12p (G) in seminomas (blue) and non-seminomas (purple), using AmplificationTimeR. Each radial line indicates the relative event ordering (T#). CNV trajectories are annotated (e.g., 5+2 GWG). The sequence of G and W denotes the chronological order of genomic events leading to the observed final copy number state. For example, “5+2 GWG” indicates that the final copy number state of 5+2 was reached through an initial gain event (G), followed by a whole-genome doubling (W), and a subsequent gain event (G) occurring after the doubling. WGD timing events are highlighted in red. Confidence intervals shown from 500 bootstraps. Only SBS1 and SBS5 mutations were used.

**Supplementary Fig. 19:** AmplificationTimeR timing analysis showing early WGD relative to 12p amplification in seminomas (blue) and non-seminomas (purple), using AmplificationTimeR. Each radial line indicates the relative event ordering (T#). CNV trajectories are annotated (e.g., 5+1 GWG). The sequence of G and W denotes the chronological order of genomic events leading to the observed final copy number state. For example, “5+2 GWG” indicates that the final copy number state of 5+2 was reached through an initial gain event (G), followed by a whole-genome doubling (W), and a subsequent gain event (G) occurring after the doubling. WGD timing events are highlighted in red. Confidence intervals shown from 500 bootstraps. Only SBS1 and SBS5 mutations were used.

**Supplementary Fig. 20:** Differential enrichment of gene sets between non-seminomas and seminomas using ssGSEA scores. (**a**) Hallmark pathways. (**b**) KEGG pathways. Circle size indicates −log₁₀(FDR); color represents relative enrichment in non-seminomas. P-values from Wilcoxon rank-sum tests; multiple testing corrected using the Benjamini–Hochberg method.

## Supplementary Data

**Supplementary Data 1:** Characterization of whole-genome sequencing quality metrics, genomic features, and clinical information for 252 TCGA TGCT tumors.

**Supplementary Data 2:** Non-synonymous somatic mutations and their inferred clonality in TCGA TGCT tumors.

**Supplementary Data 3:** Somatic copy number analysis of chromosome X in TCGA TGCT tumors, generated using the Battenberg algorithm.

**Supplementary Data 4:** Estimated replication stress scores (RepStress_ZScore) for all TCGA tumor samples.

**Supplementary Data 5:** Mutational signature activities (SBS, ID and DBS) for TCGA TGCT tumors.

**Supplementary Data 6**: Telomere length (TL) estimates for TCGA TGCT tumors, estimated using the Telseq algorithm.

**Supplementary Data 7:** Clonality analysis of mutational clusters identified in TCGA TGCT tumors.

**Supplementary Data 8:** Identification and timing analysis of whole-genome doubling events in TCGA TGCT tumors.

**Supplementary Data 9**: Single-sample gene set enrichment analysis of TCGA tumors using Hallmark gene sets derived from RNA-seq data.

**Supplementary Data 10**: Single-sample gene set enrichment analysis of TCGA tumors using KEGG gene sets derived from RNA-seq data.

