## Supplementary Figures for "A Thiopurine-like Mutagenic Process Defines TGCT Subtypes"

**Supplementary Fig. 1**

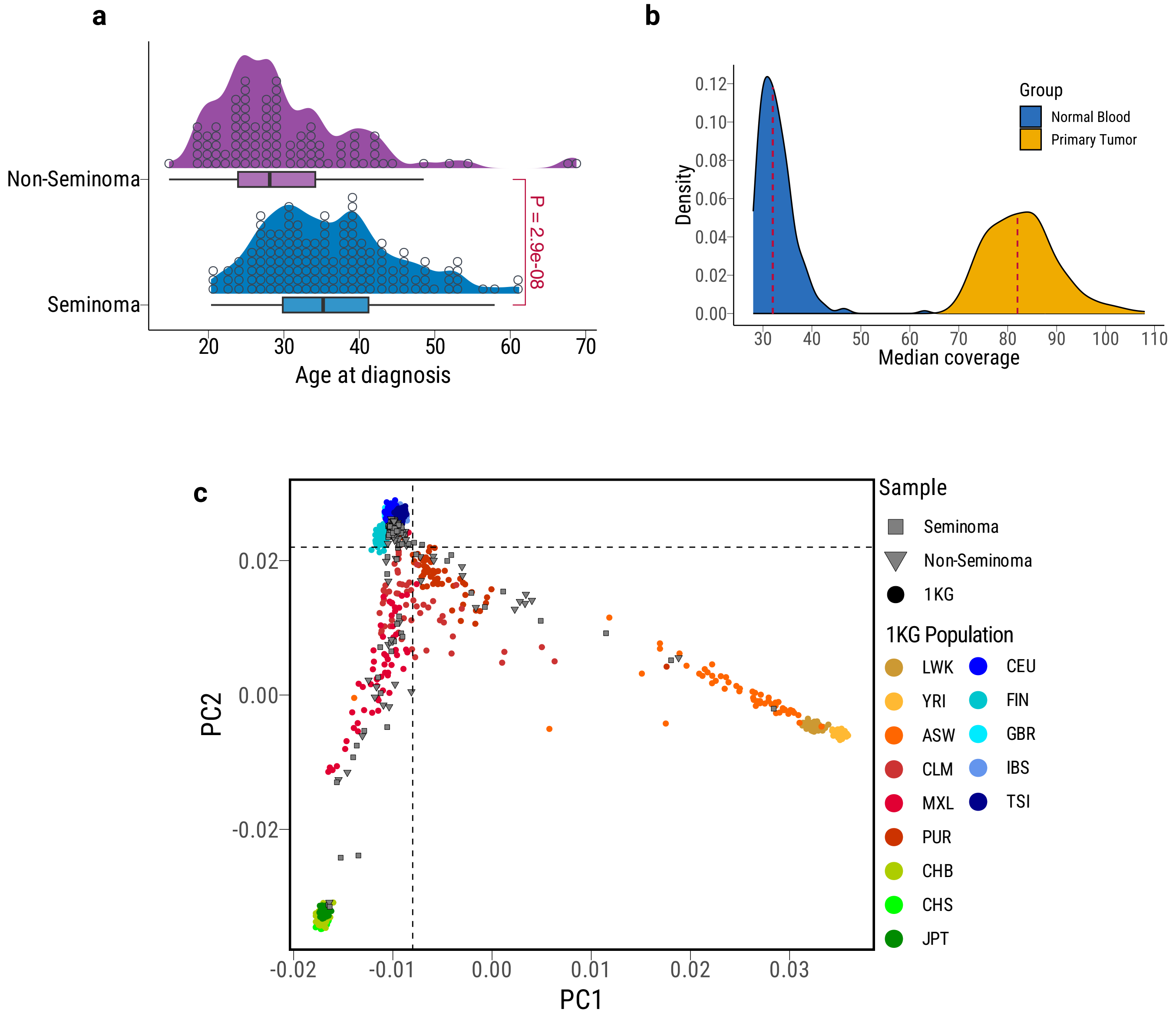

**Supplementary Fig. 2**

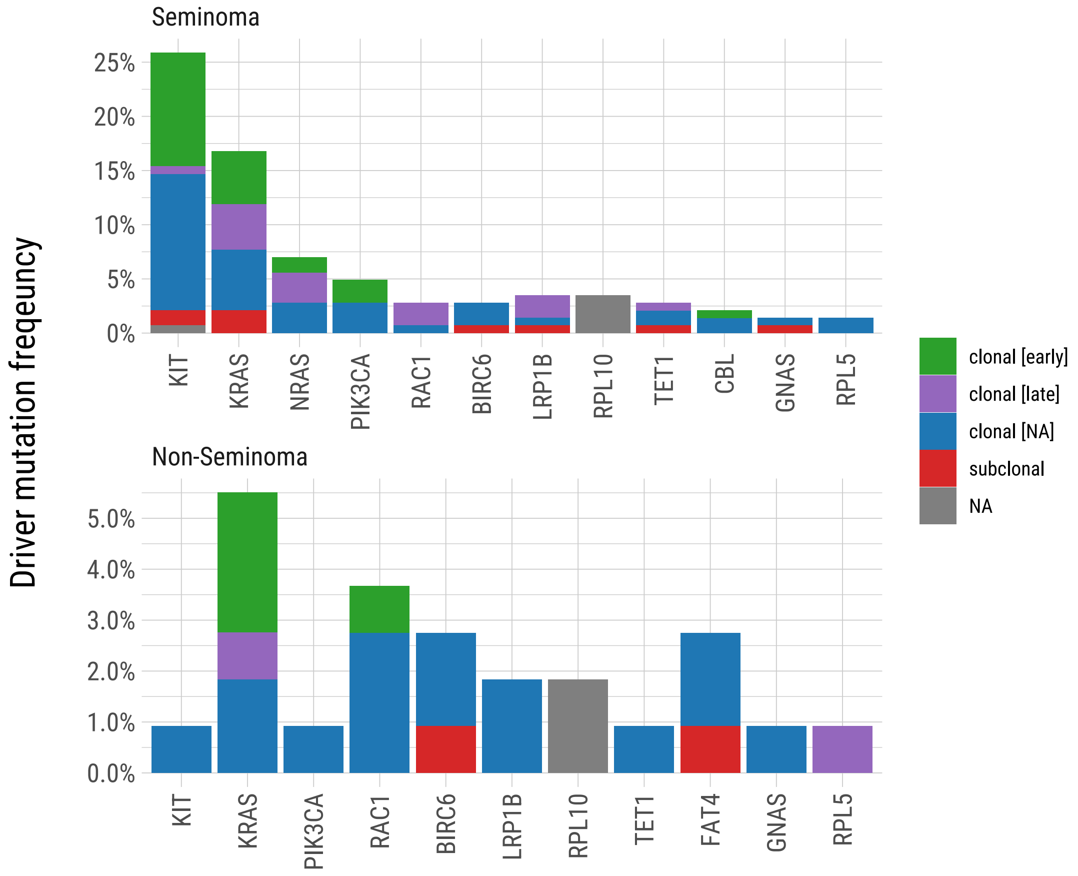

#### Supplementary Fig. 3

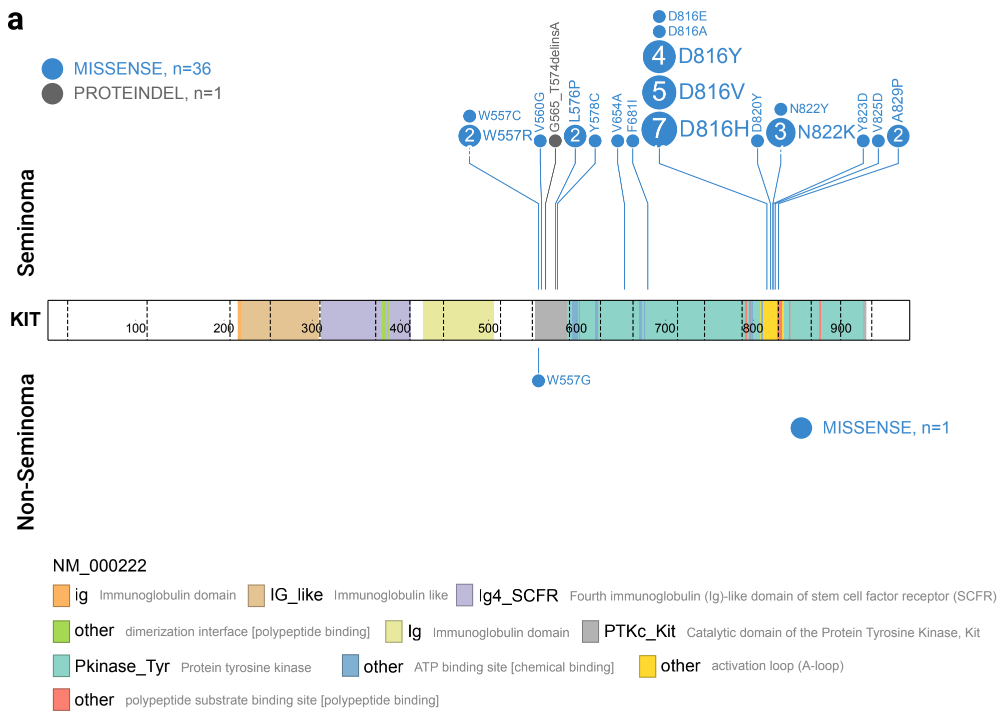


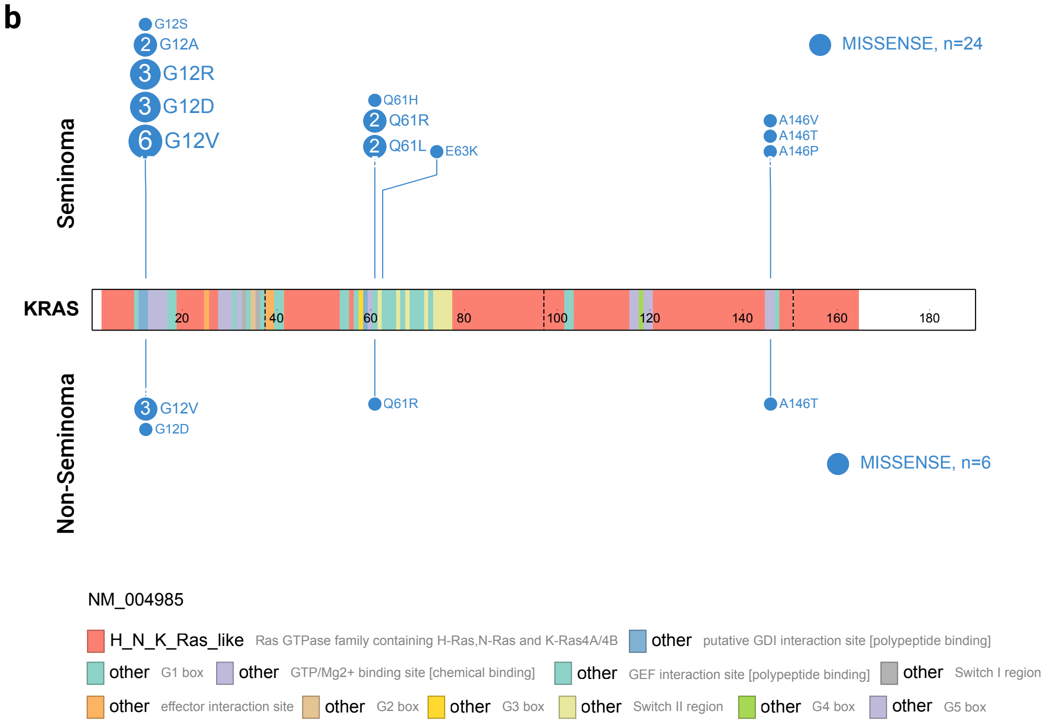

#### Supplementary Fig. 4

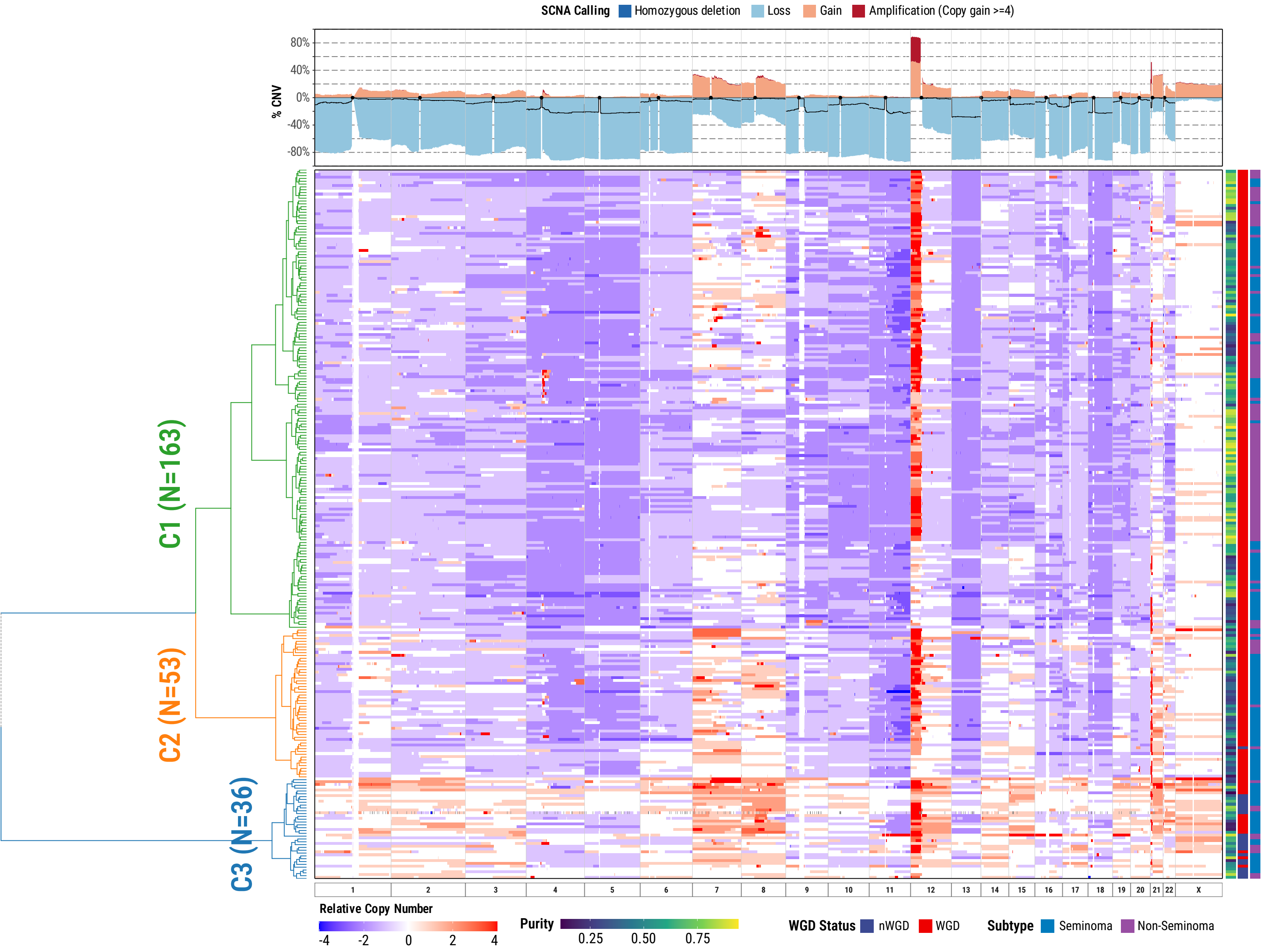

### Supplementary Fig. 5

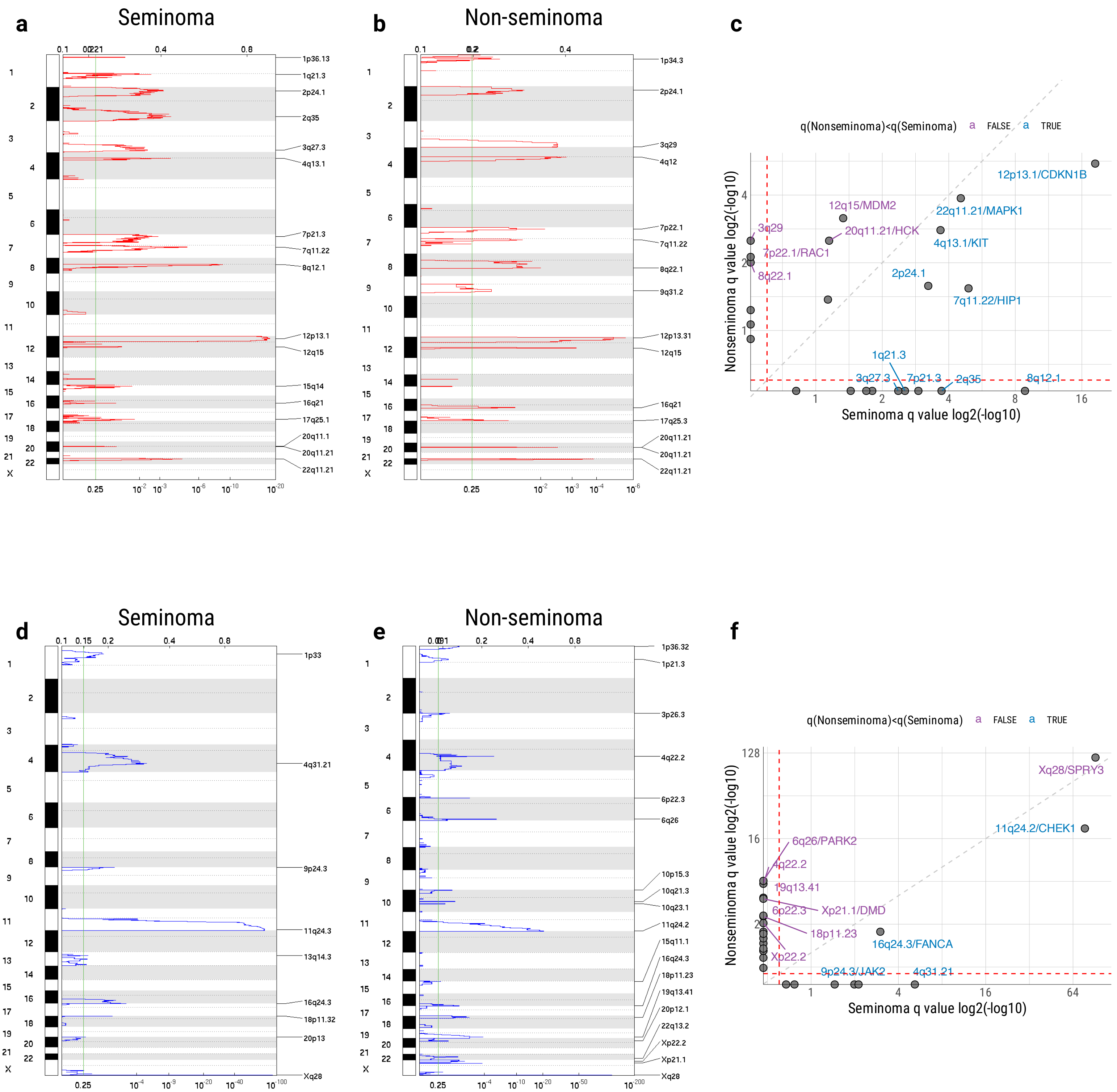

#### Supplementary Fig. 6

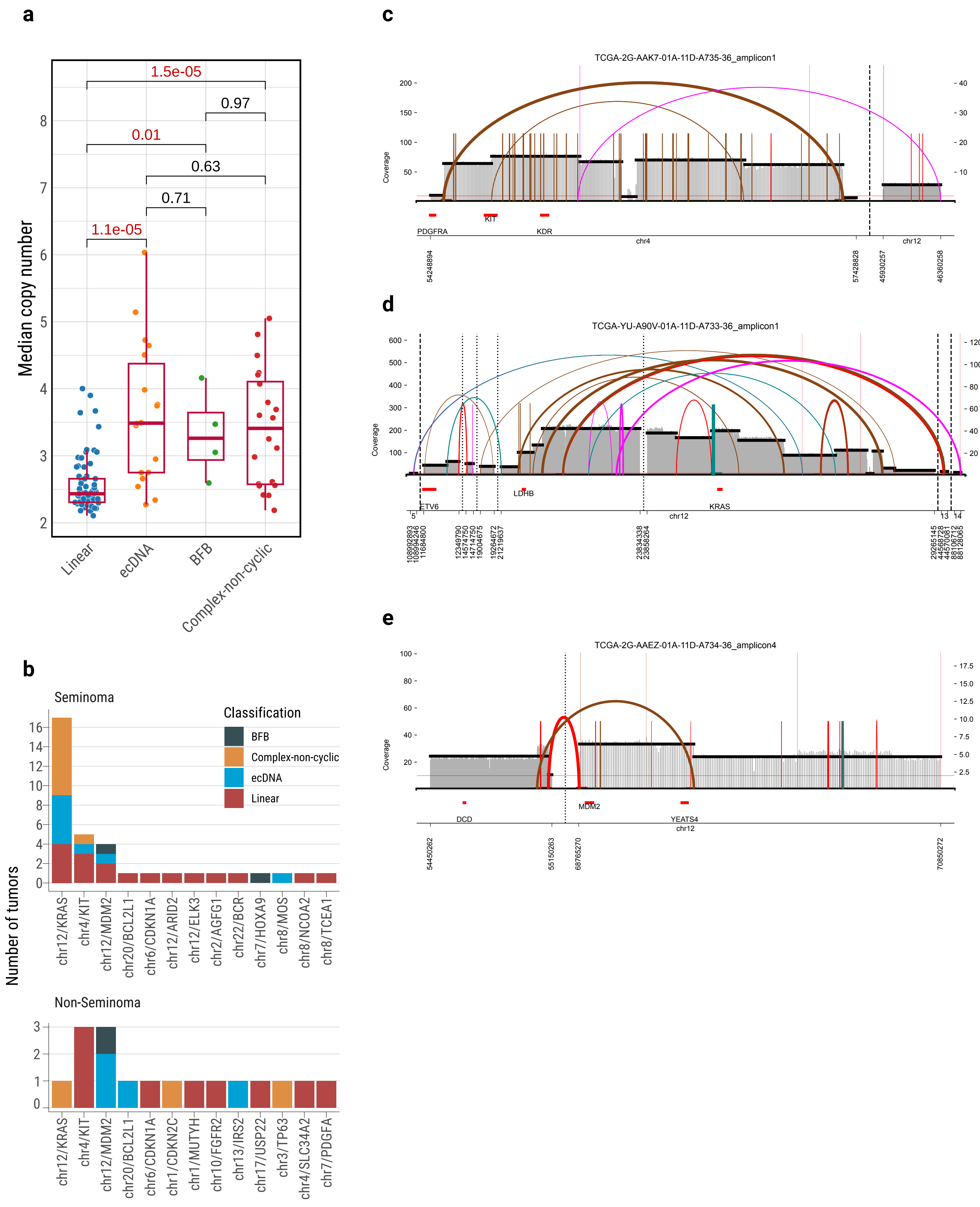

**Supplementary Fig. 7**

**a**

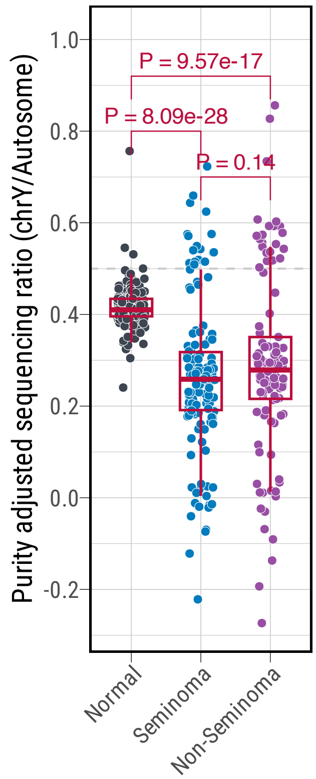


**b**

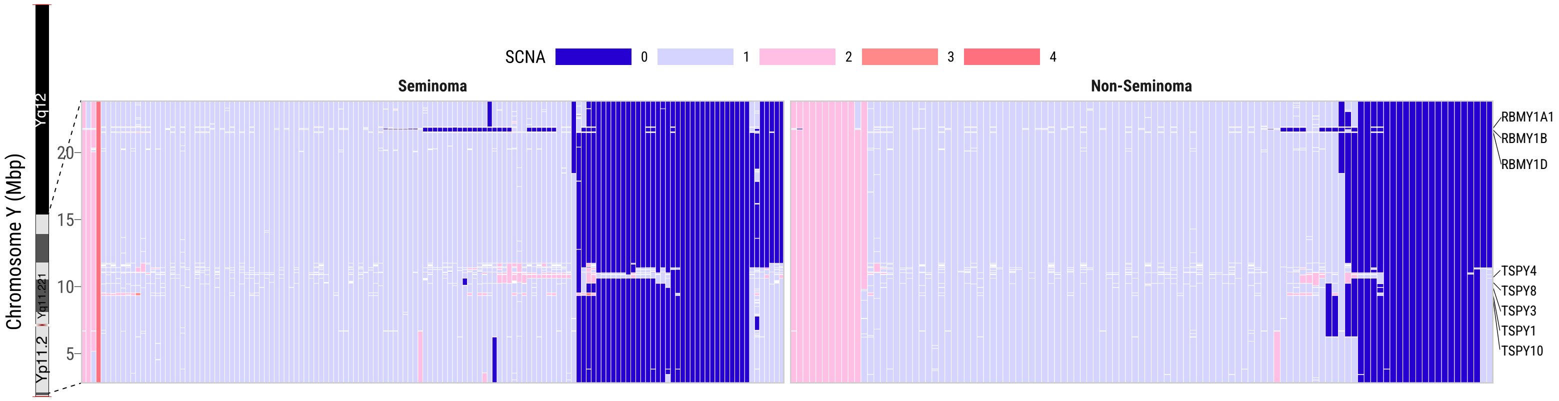

**Supplementary Fig. 8**

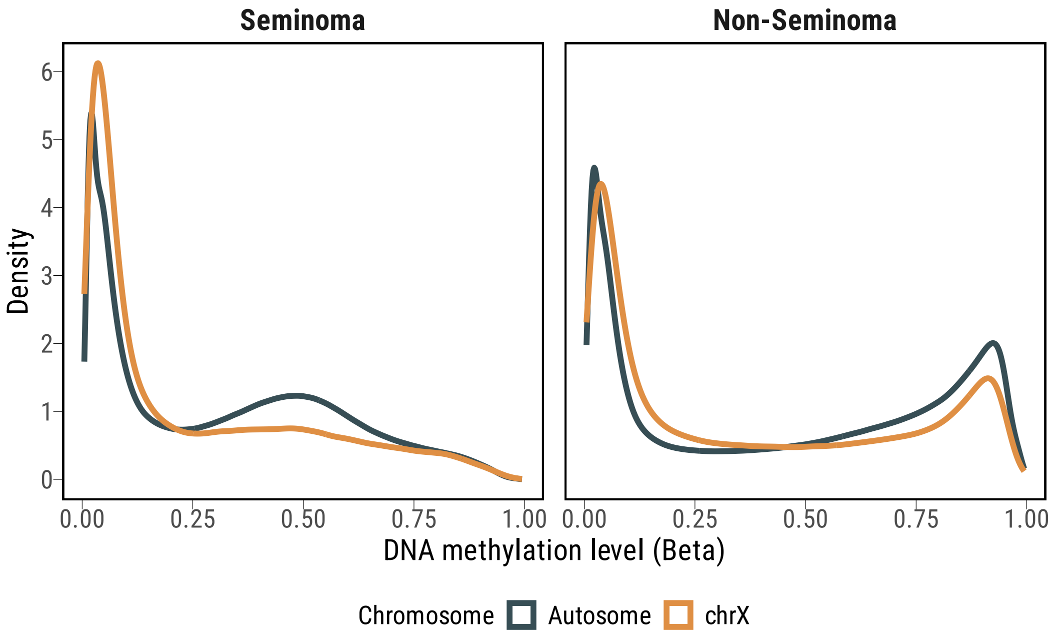

**Supplementary Fig. 9**

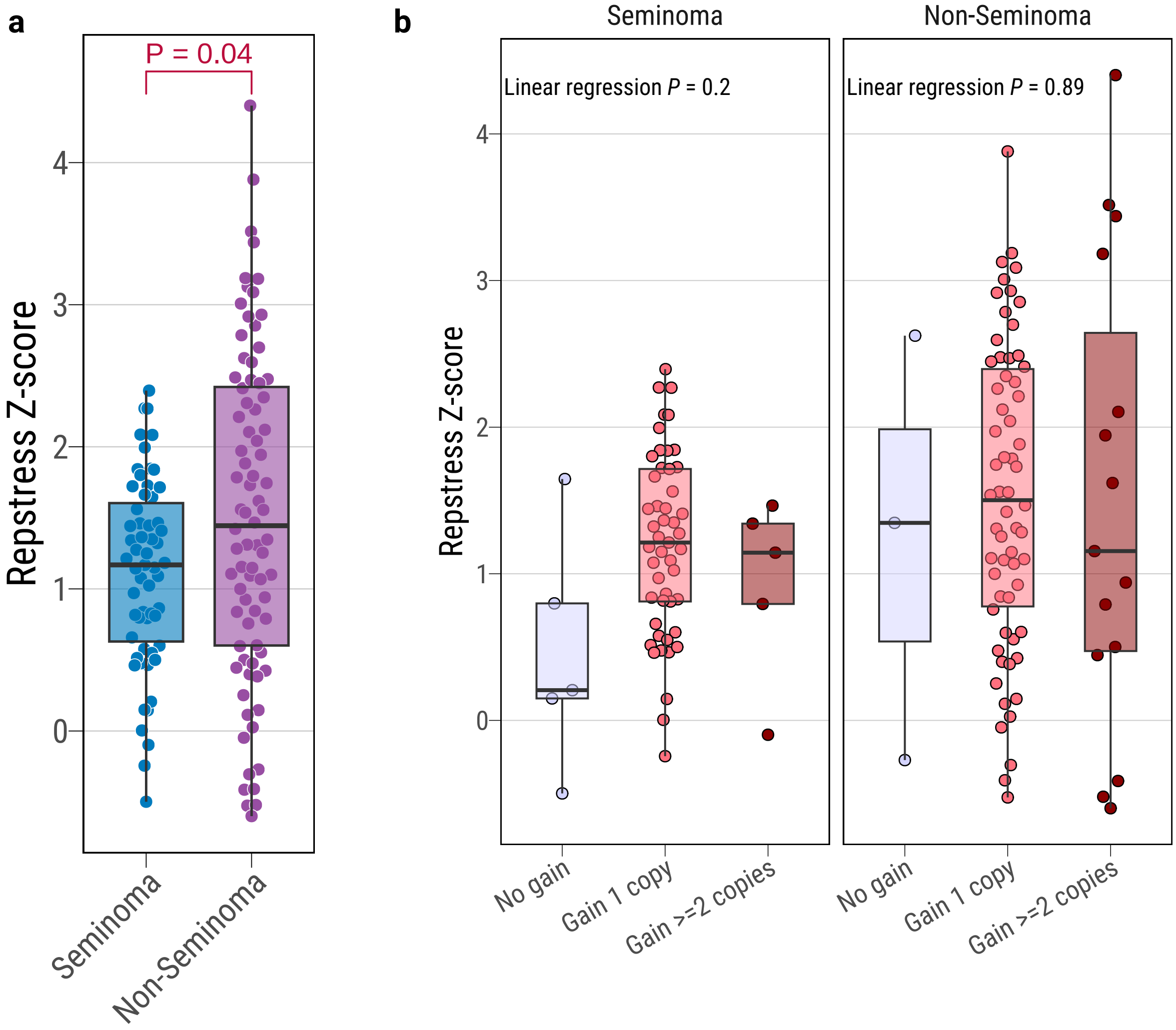

**Supplementary Fig. 10**

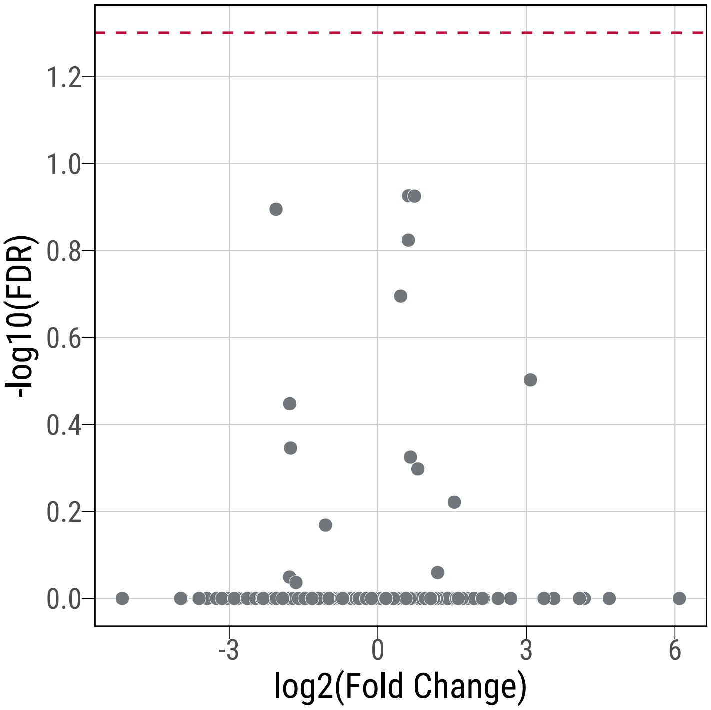

**Supplementary Fig. 11**

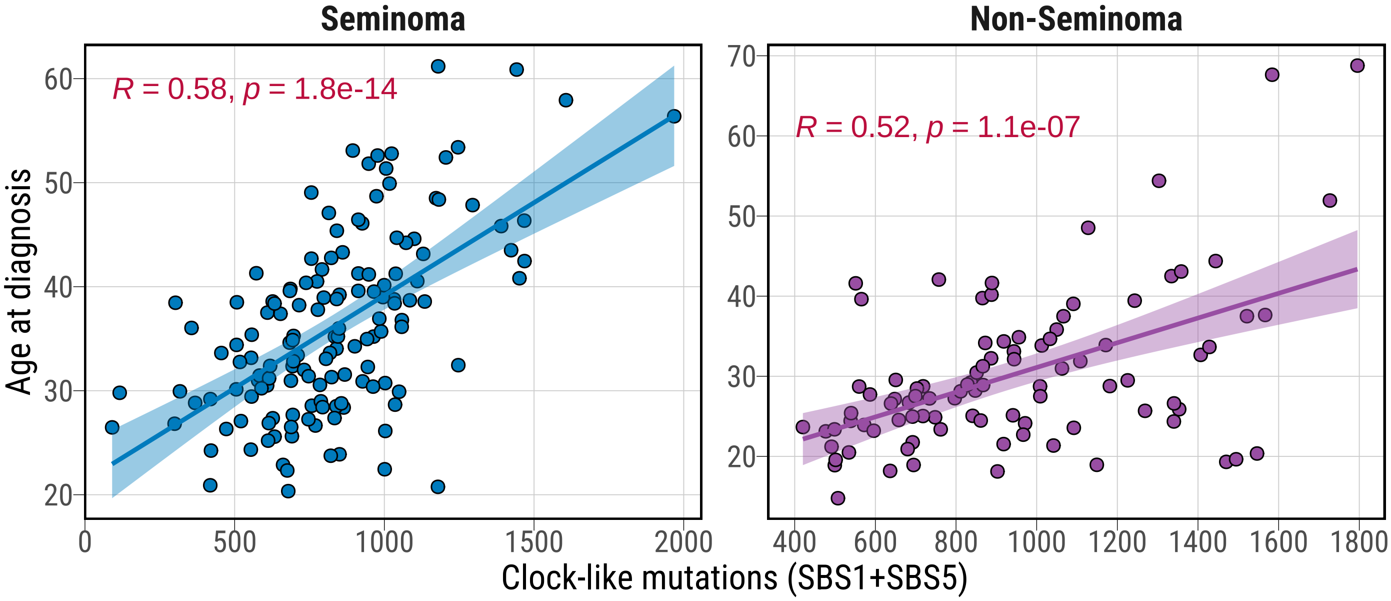

Supplementary Fig. 12

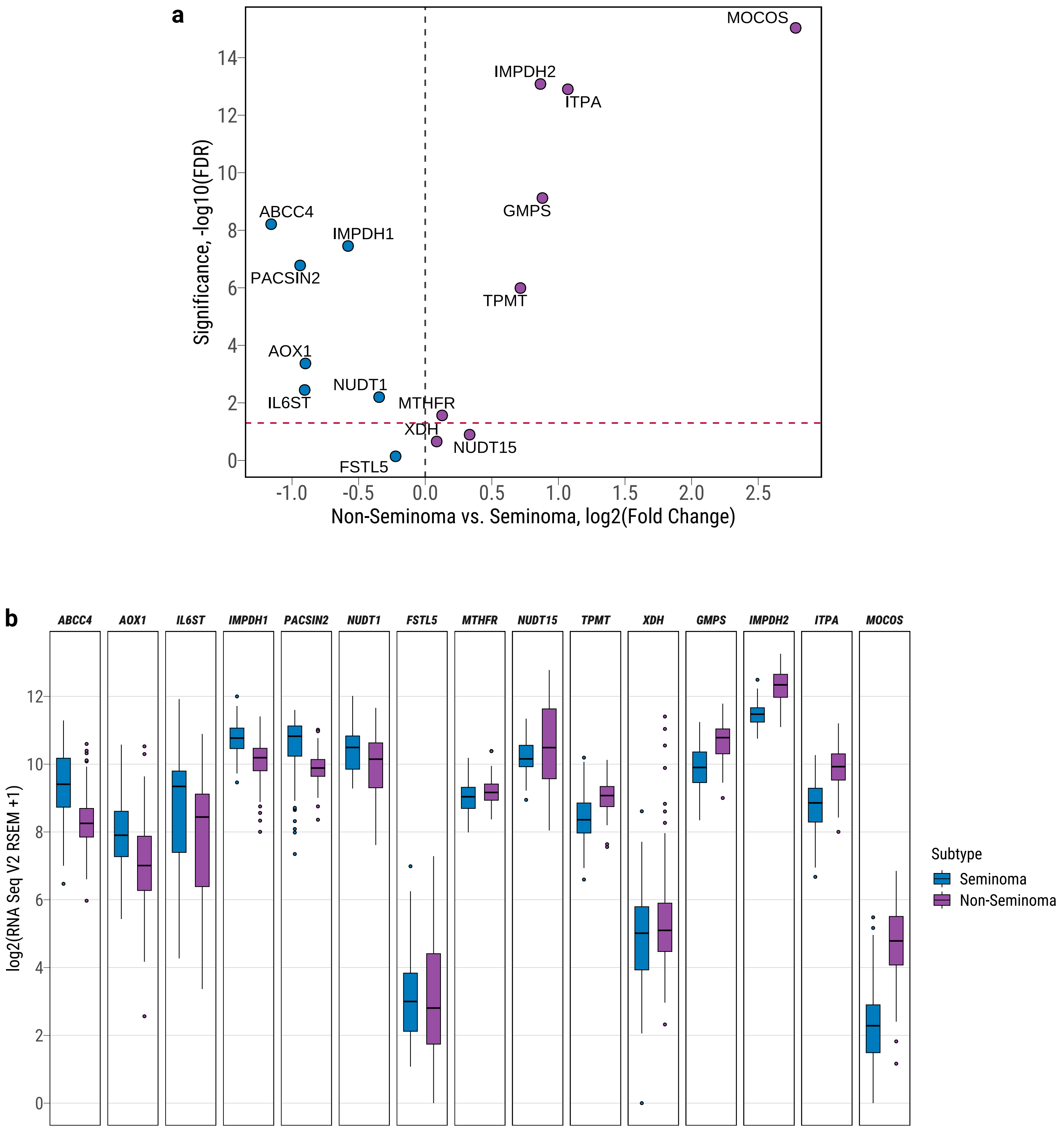

**Supplementary Fig. 13**

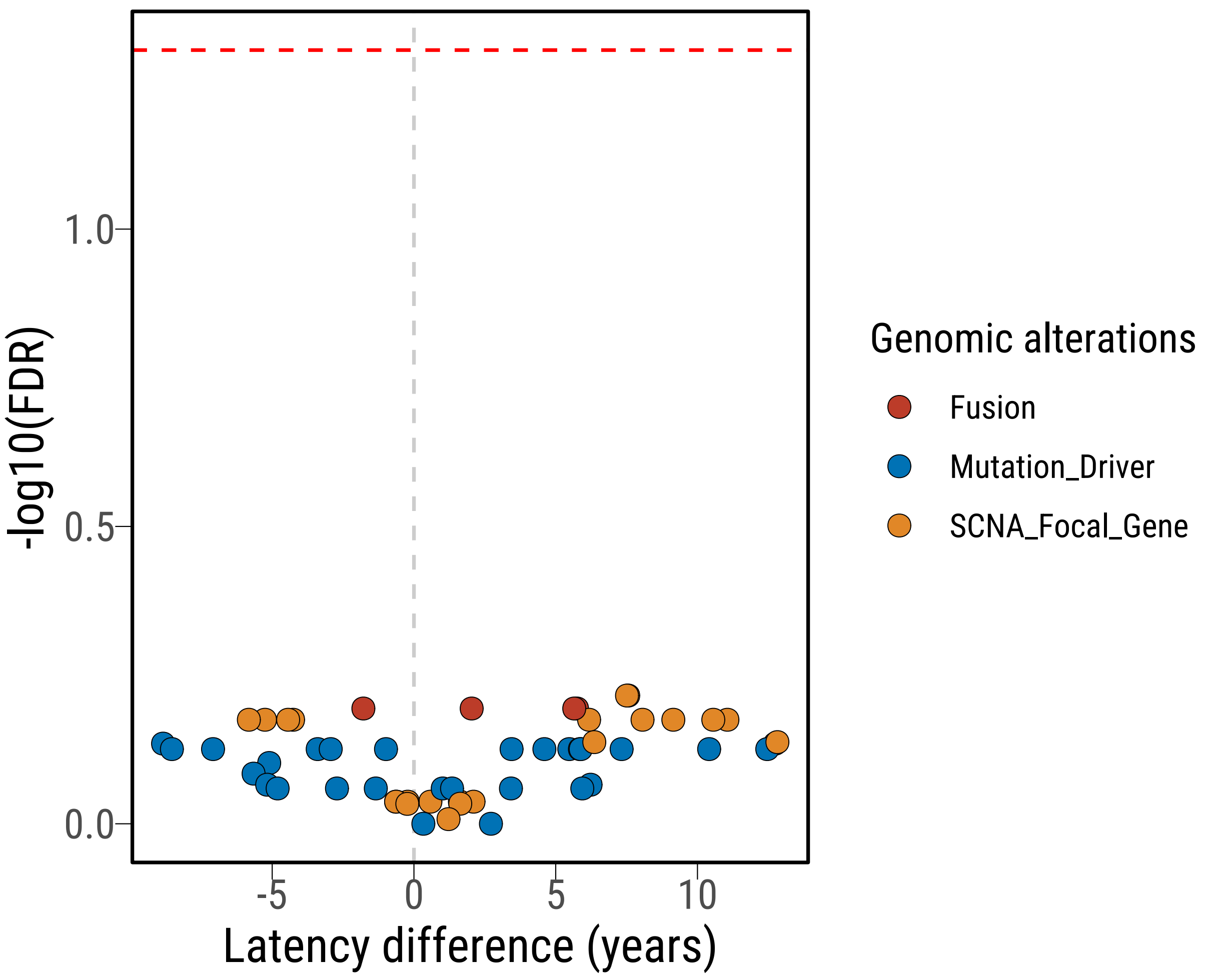

**Supplementary Fig. 14**

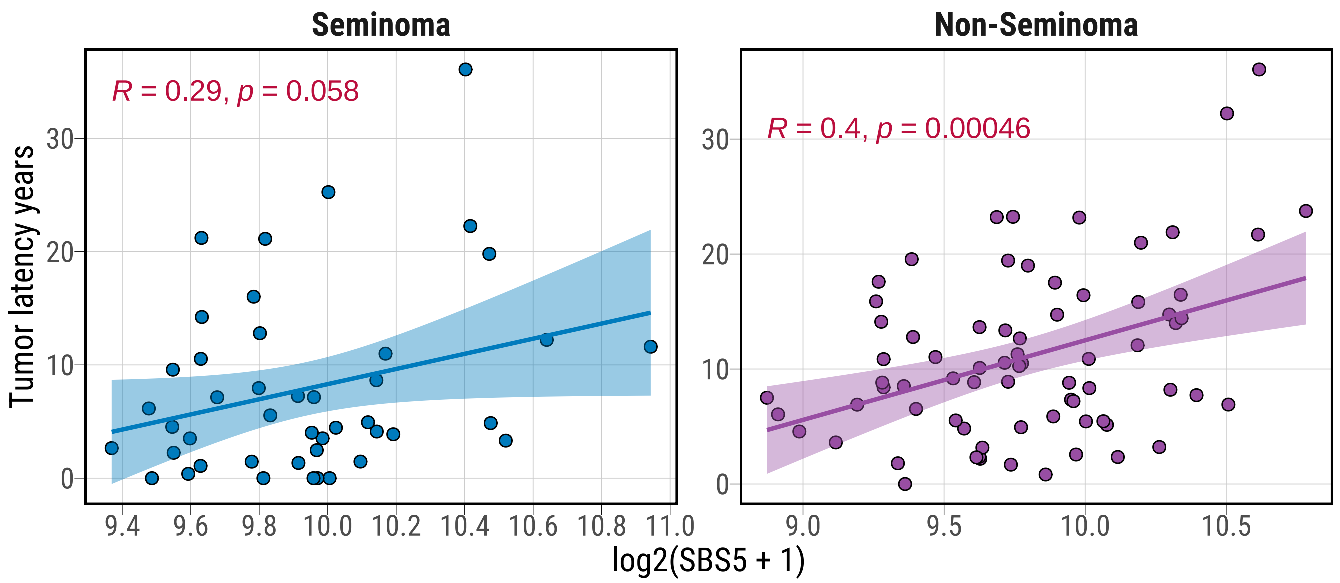

**Supplementary Fig. 15**

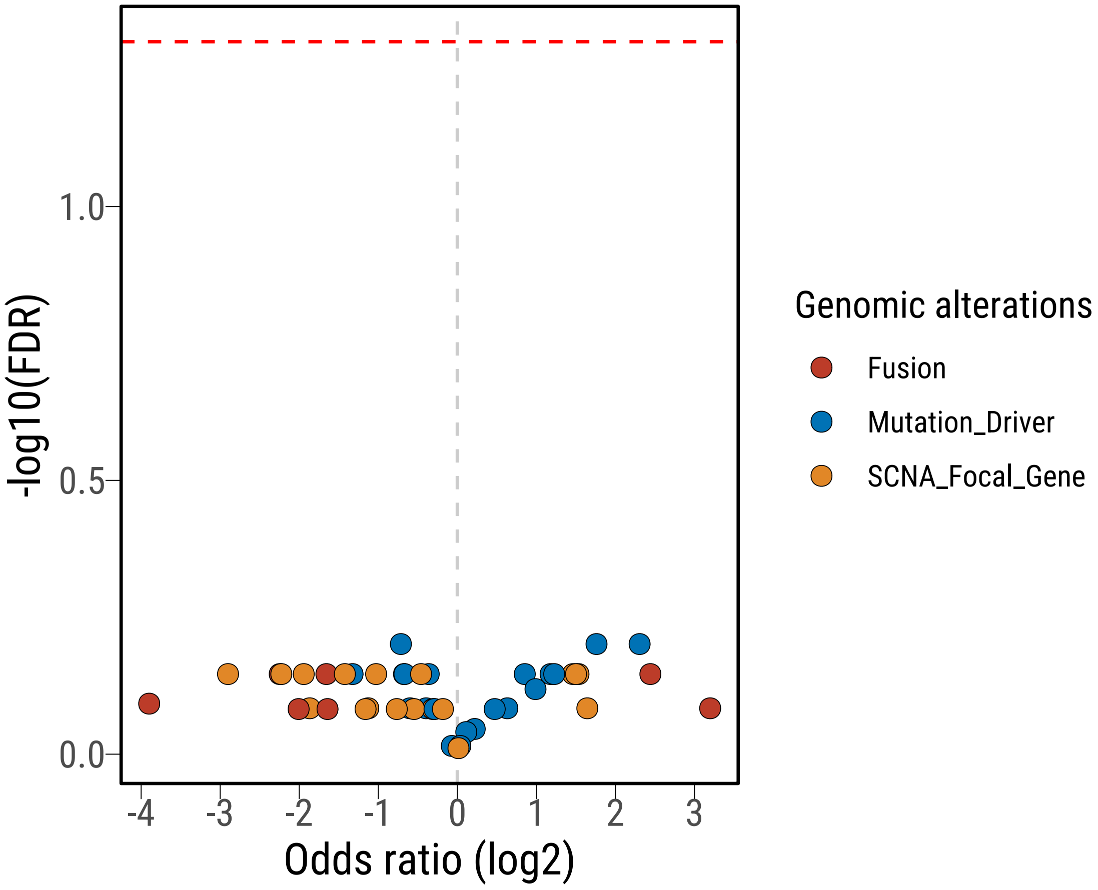

**Supplementary Fig. 16**

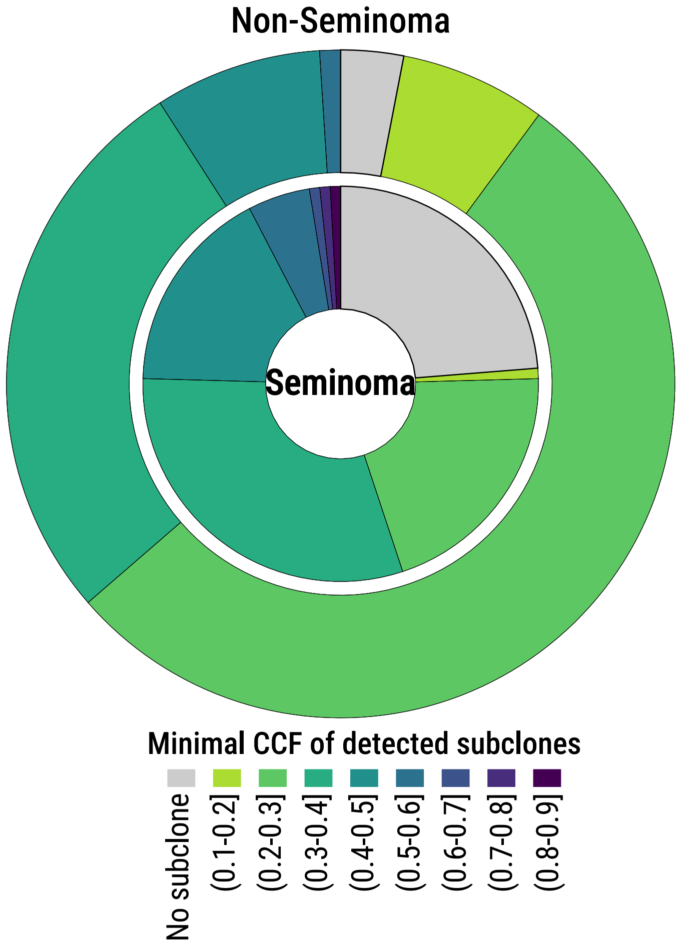

**Supplementary Fig. 17**

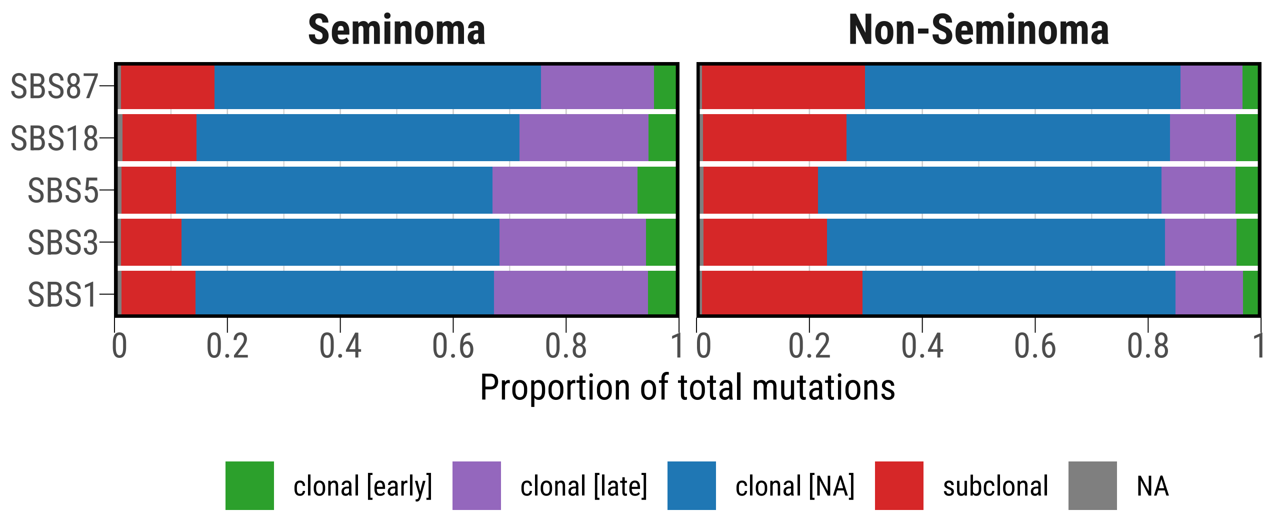

**Supplementary Fig. 18**

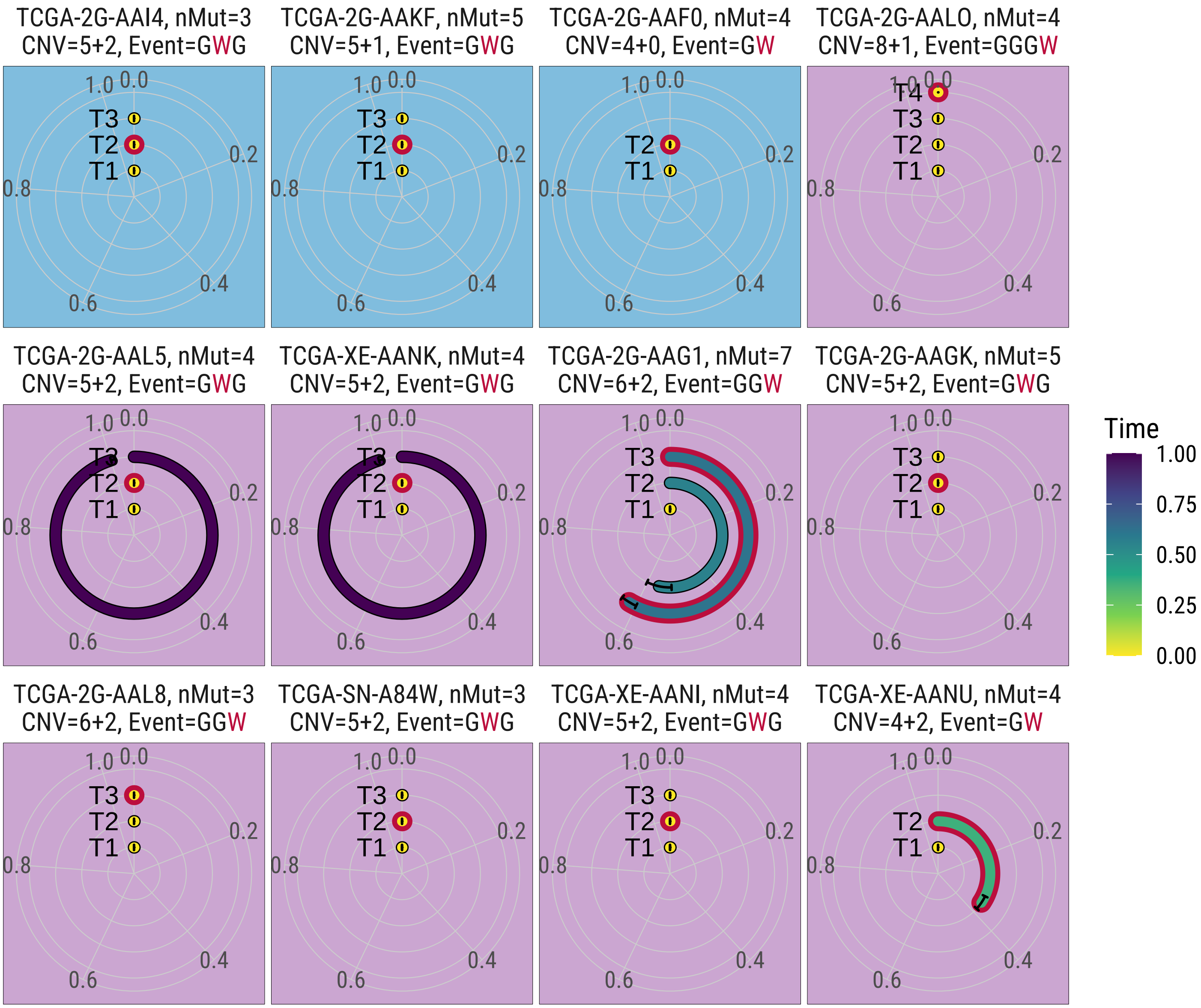

### Supplementary Fig. 19

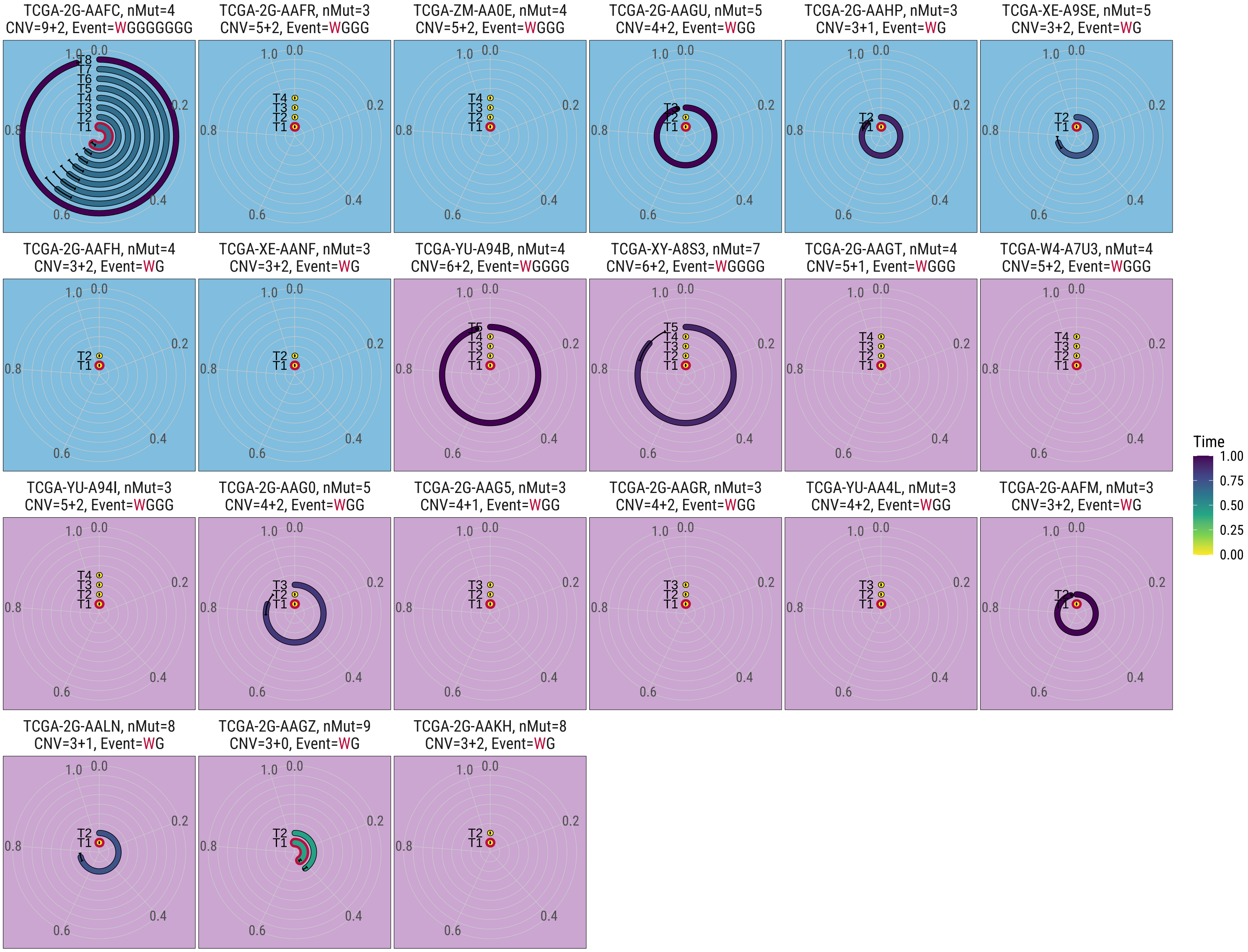

### Supplementary Fig. 20

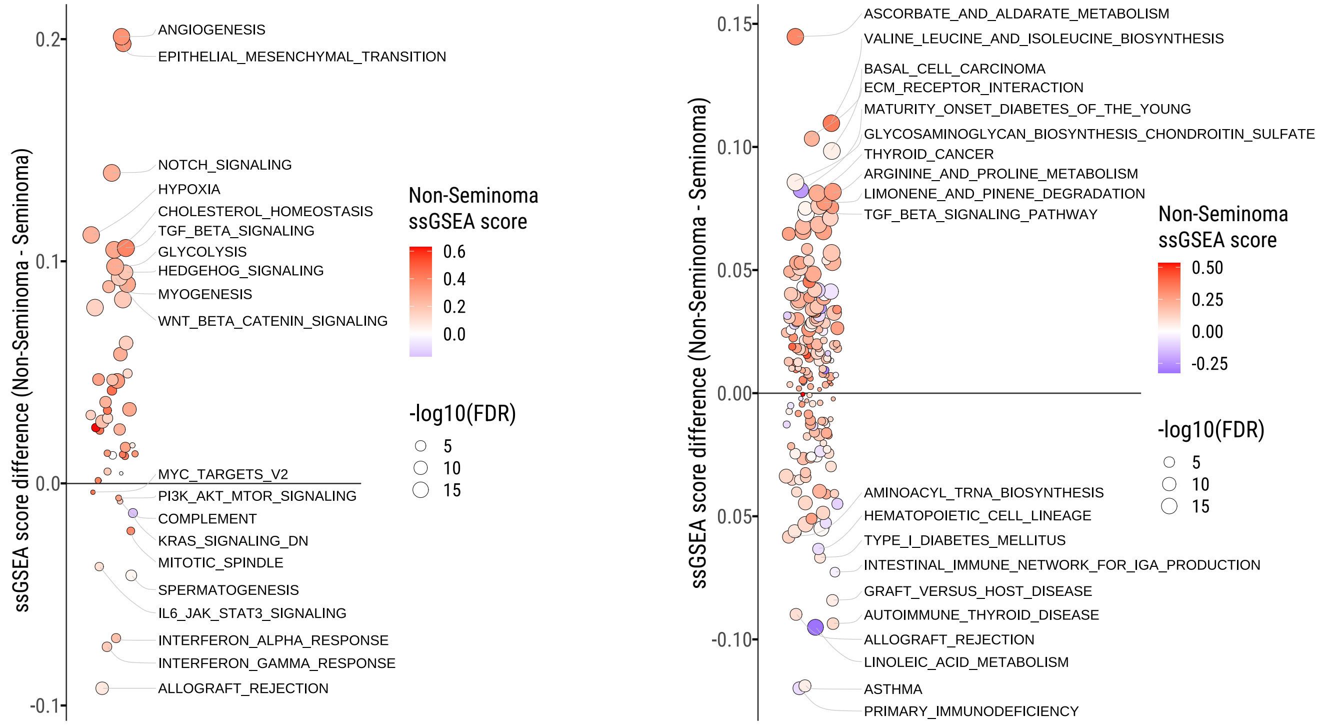
